# Functional Dynamics Reveal the Response of the Crabapple (*Malus* sp.) Phyllosphere Microbiome to *Gymnosporangium yamadae* Infection

**DOI:** 10.1101/2025.02.11.637274

**Authors:** Qianran Xu, Yonghua Zhang, Siqi Tao

## Abstract

The plant microbiome is crucial for maintaining plant health under disease stress. While considerable research has focused on belowground systems, the phyllosphere microbiome remains underexplored, particularly regarding its functional responses to pathogen infection. In this study, we investigated the phyllosphere microbiome dynamics of crabapple (*Malus* ‘Kelsey’) infected by *Gymnosporangium yamadae* using metatranscriptomic sequencing across six stages of rust disease progression. Our analysis revealed a general increase in the diversity of fungi, bacteria and viruses in infected leaves. Notably, fungal diversity negatively correlated with bacterial diversity, reflecting competitive interactions during disease development. Microbial taxa in diseased leaves exhibited heightened expression activity compared to healthy leaves, with fungi progressively dominating the microbial community. Functional co-occurrence networks of the phyllosphere microbiome in infected leaves were more complex than in healthy leaves, suggesting adaptive reorganization in response to pathogen invasion. Differentially expressed genes at each stage were significantly enriched in carbohydrate metabolism pathways and gene regulation-related functions, enabling functional adaptations to rust diseases. Random forest modeling identified key microbial transcripts associated with pathogen abundance, including beneficial microbes like *Saitozyma podzolica*, which secretes glucan-degrading enzymes, and *Mortierella elongata*, involved in sterol biosynthesis and plant resistance. Conversely, *Alternaria alternata* emerged as a major pathobiome contributor, secreting enzymes that degrade plant cell wall components (e.g., pectin, cellulose, lignin, and xylan), and engaging in MAPK signaling pathways critical for pathogenesis. Our findings underscore the vital role of the phyllosphere microbiome in mediating plant-pathogen interactions and shaping disease progression, providing a foundation for microbiome-based strategies to enhance plant resilience.

**Importance:** Our findings reveal the dynamic shifts in the expression patterns and adaptive functional strategies of crabapple phyllosphere microbiome in response to the *Gymnosporangium yamadae* infection. We identified several key microbes that may play vital roles in pathogenesis and speculated on their roles and functions in plant-pathogen interactions. In conclusion, our study highlights the potential of the phyllosphere microbiome in regulating plant health.

## 1 Introduction

The plant microbiome encompasses an entire community of microorganisms and their associated molecular spectrum within plant environments, forming a diverse and complex ecological network that significantly impacts plant health, resilience and defense mechanisms ^[1–5]^. Current research underscores that plant fitness and immunity are not shaped solely by individual pathogens or beneficial species but by the collective interactions within the microbiome ^[6]^.

Beneficial microbial communities, especially within the rhizosphere and phyllosphere, play critical roles by modulating immune responses, competing with pathogens, and producing metabolites that enhance defense ^[3,7–9]^. For instance, *Pseudomonas* and *Bacillus* in the rhizosphere produce antibiotics that directly suppress pathogens ^[10,11]^. Furthermore, plants can actively engage in recruiting beneficial microbes, particularly under pathogen pressure, to alleviate stress and restructure microbial communities assemblies to enhance resilience ^[12,13]^. For example, the rhizosphere microbiome of the *Ralstonia solanacearum*-resistant tomato variety Hawaii 7996 has a higher abundance of beneficial microbes, such as *Flavobacteriaceae*, *Sphingomonadaceae* and *Pseudomonadaceae* compared to the susceptible variety Monkeymaker, providing a natural microbial buffer against pathogens ^[14]^. Similarly, *Arabidopsis thaliana* recruits protective bacteria, such as *Microbacterium*, *Stenotrophomonas*, and *Xanthomonas*, which collectively induce systemic immunity against downy mildew ^[15]^. Likewise, in wheat, exposure to high pathogen pressure from *Fusarium pseudograminearum* leads to an enrichment of *Stenotrophomonas rhizophila*, functioning as an early defense alert in the root endosphere ^[16]^. This observed increase in protective microbial abundance in various plant-pathogen systems aligns with the studies showing that stressed plants under pathogen pressure can engage in a “cry for help”, attracting beneficial microbes that bolster plant defenses ^[15,17,18]^.

As research advances, scientists increasingly recognize the potential of leveraging microbiome functions for sustainable crop protection and yield improvement ^[3]^. Functional studies have linked specific gene expression profiles to pathogenic responses, further deepening our understanding of microbiome interactions ^[19]^. For example, during *Fusarium* wilt in peppers, genes associated with detoxification, biofilm formation, and chemotaxis were significantly enriched, suggesting targeted adaptations in the root endosphere, a coordinated microbial response to bolster plant defenses ^[20]^. Similarly, *Rhizoctonia solani* infection in sugar beets led to an increase in *Chitinophagaceae* and *Flavobacteriaceae*, which produced antifungal enzymes and metabolites like phenazines, polyketides, and siderophores, providing an additional line of defense against pathogens ^[21]^.

However, not all microbial members benefit plant health, some can be detrimental by forming harmful partnerships with pathogens, disrupting plant resilience, and facilitating disease progression ^[6,22,23]^. For example, *Verticillium dahlia* infection was shown to stabilize certain microbial networks within the bulk soil and the rhizosphere, with viruses becoming as central players in the ‘pathobiome’, potentially aiding pathogen invasion and colonization ^[24]^. Similarly, studies on the diseases affecting tomato, tobacco and other plants have shown that certain microbes associated with nematode pathogens release enzymes that degrade plant cell walls, which aids pathogen entry into roots and exacerbates disease progression ^[25,26]^. This multifaceted relationship underscores the role of the plant microbiome in pathogenesis cannot be viewed as a straightforward dichotomy of pathogenic or protective effects, but must be analyzed in context ^[27]^.

Despite the insights gained from studies on the rhizosphere, most research on plant-microbiome-pathogen interactions has focused on the belowground microbiome, particularly the rhizosphere. In contrast, the aboveground microbiome, or phyllosphere-constituting the above ground plant microbiome-remains relatively underexplored despite it holds significant potential as a critical line of defense against airborne pathogens ^[28–30]^. Under abiotic stress, the *Arabidopsis* mutant *mfec* exhibited significantly reduced phyllosphere microbiome diversity, which was linked to the loss of immune and MIN7 pathways, resulting in dysbiosis and plant symptoms. This underscores the critical role of phyllosphere microbiome homeostasis in maintaining plant health ^[29]^. Increasing evidence also suggests the assembly strategies of the phyllosphere microbiome are dynamic, with strategies varying in response to different pathogen pressures ^[31]^. For instance, the phyllosphere microbiome of citrus plants infected with *Diaporthe citri* exhibited lower community evenness and a more complex co-occurrence network, with the emergence of new microbial taxa ^[30]^. Similarly, temporal analysis of the phyllosphere and rhizosphere microbiomes in *Phytophthora sojae*-infected and *Septoria glycines*-infected soybean plants revealed that the structure of the phyllosphere microbial communities was more responsive to pathogen infection and disease progression, with a noticeable increase in the abundance of saprophytic fungi ^[32]^. During disease progression, the phyllosphere microbial communities in apple, wheat and tobacco were observed to exhibit higher diversity and more complex co-occurrence networks ^[8,33–35]^. However, while many of the studies to date has focused on identifying the microbial components of the disease-induced phyllosphere microbiome, the functional responses of these communities and their direct links to plant health have been largely overlooked ^[36]^.

Crabapple trees (*Malus* spp.), cherished in landscaping for their attractive shape and leaf color, face a significant threat from the rust fungus *Gymnosporangium yamadae*, which severely diminishes their ornamental value ^[37]^. *G. yamadae*, a heteroecious fungal pathogen, requires two different hosts (*Malus* spp. and *Juniperus chinensis*) to complete its infection cycle and produces four morphologically distinct spores ^[38,39]^. In early spring, brownish telia break through the epidermis of the teilal host (*J. chinensis*), forming a bright yellow gelatinous mass and releasing haploid basidiospores that infect *Malus* species. Initial infection manifests as chlorotic spots and the development of yellowish droplets (spermogonia) on the upper surface of the infected leaves ^[38]^. The biotrophic and unculturable nature of *G. yamadae* complicates traditional pathogen-host studies, prompting researchers to pivot towards investigating the response patterns of microbiome ^[13]^. Recent studies have highlighted the essential role of microbiome in modulating plant responses to biotrophic pathogens. For example, studies of the phyllosphere microbiome in cucumber infected with powdery mildew cucumber revealed significant differences in bacterial alpha diversity across varying disease severity, with more severely diseased leaves exhibiting higher bacterial diversity and an increased abundance of certain beneficial microbes ^[40]^. Similarly, in *Arabidopsis thaliana* infected with the downy mildew pathogen *Hyaloperonospora arabidopsidis*, three bacterial species were specifically enriched in the rhizosphere, collectively inducing systemic resistance and promoting plant growth ^[15]^. In the studies of apple and *G. yamadae* biotrophic interactions, a meta-transcriptome analysis was conducted to compare the phyllosphere fungal communities in infected and uninfected apple leaves (*M. domestica* cv. Fuji) at 10- and 30-days post-inoculation (dpi). This analysis revealed a significant shift in the community composition occurs during the later stages of infection, with a notable increase in the abundance of *Alternaria* and *Fonsecaea* species at 30 dpi ^[34]^. Subsequent research expanded on these findings with a comprehensive time-course analysis of fungal and bacterial community dynamics, as well as key leaf metabolites, in two crabapple varieties, ‘Flame’ and ‘Kelsey’, over six distinct stages of rust disease progression ^[8]^. The integrative study highlighted that the leaves regulate disease progression by secreting specific metabolites, which help mediate the enrichment of potential beneficial microbes, supporting the “cry for help” strategy, where plants under stress recruit microbial allies for defense ^[8]^. A complementary investigation explored the diversity and structural dynamics of endophytic microbial communities in apple (*Malus domestica* cv. Gala) leaves across various stages of rust infection. This study utilized amplicon sequencing to offer foundational insights into the predicted functional profiles of endophytic fungi and bacteria, shedding light on the microbial shifts associated with disease progression ^[41]^. While these studies have advanced our understanding of the diversity and structural changes in the phyllosphere microbiome of *Malus* species under *G. yamadae* infection, a comprehensive understanding of the dynamic functional profiles of these microbial communities throughout the disease course is still lacking. Such insights are critical for uncovering the adaptive responses of *Malus* phyllosphere microbiome to *G. yamadae* infection and understanding their impact on host plant health.

This study aims to systematically investigate the functional response of the phyllosphere microbiome in crabapple leaves infected by *G. yamadae* at various stages of lesion expansion using meta-transcriptomic technology. Our main objectives were to (i) assess the diversity and composition of phyllosphere microbial transcriptomes at various stages of lesion expansion; (ii) pinpoint differentially expressed microbial genes at each stage of infection and examine their dynamic expression profiles over time; (iii) elucidate the functional roles and activities of phyllosphere microbiota and evaluate their potential interactions with plant disease processes and plant health.

## 2 Results

### 2.1 The taxa transcriptomes diversity and structure patterns in the phyllosphere with infection

The phyllosphere of crabapple leaves harbors a complex microbial community whose diversity and structure undergo significant shifts in response to *G. yamadae* infection. Based on the analysis of transcript expression levels and taxonomic annotations, the microbial community in the phyllosphere was categorized into bacterial, fungal, viral, and archaeal transcriptomes, with species-level expression quantified as FPKM values (Table S1). Archaea were excluded from further transcriptome diversity and structural analyses due to their low diversity.

The alpha diversity analysis of the phyllosphere microbiome during *G. yamadae* infection revealed distinct trends in bacterial, fungal, and viral transcriptomes. Overall, fungal transcriptomes consistently exhibited greater alpha diversity than that of bacterial and viral transcriptomes (Figure 1a, S1a). For bacterial transcriptomes, diseased leaves generally exhibited higher species richness compared to healthy leaves. The Shannon index in healthy leaves peaked at the third stage and declined to its lowest value by the sixth stage. In early to mid-disease stages (Stages 1 to 4), healthy leaves displayed greater bacterial diversity than diseased leaves. However, as the lesions expanded, the Shannon index and Pielou’s evenness index of bacterial transcriptomes in diseased leaves incrementally increased, peaking at the sixth stage (Figure 1a, S1b). For fungal transcriptomes, healthy leaves did not display any distinct patterns in diversity indices. However, *G. yamadae* infection initially resulted in a decline in species richness, which then rose to peak at the final stage. Interestingly, the Shannon index, as well as the Pielou’s evenness index, declined steadily in diseased leaves, an opposite trend compared to bacterial diversity (Figure 1, S1b; Table S2a). The correlation analysis between the Shannon indices of bacterial and fungal communities through the simple linear model confirmed the inverse relationship between the Shannon indices of bacterial and fungal communities, with bacterial diversity increasing the fungal diversity decreasing in diseased leaves (Figure 1b). The viral alpha diversity did not exhibit any significant patterns across stages or between leaf conditions (Figure S1a; Table S2a).

**Figure 1.**
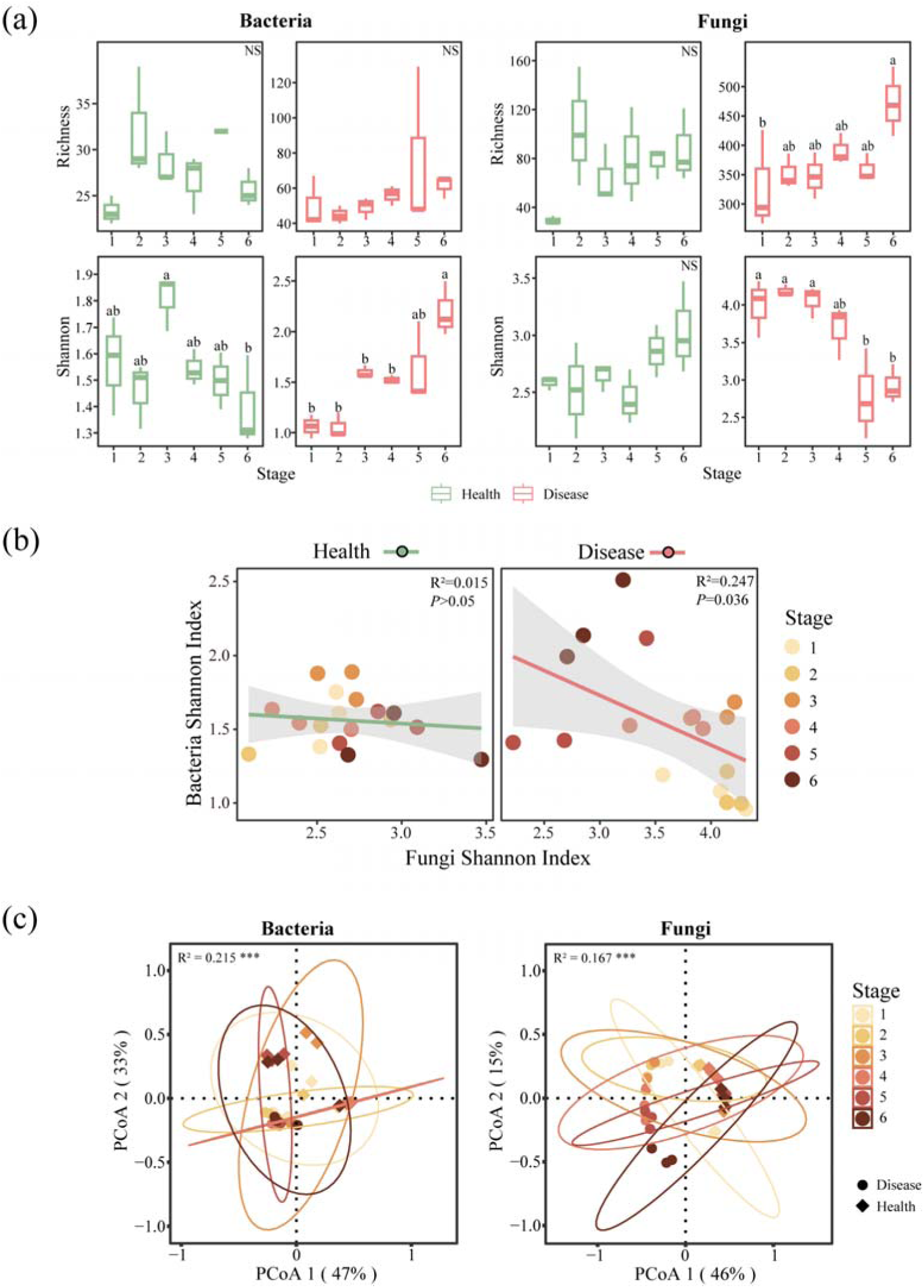
Phyllosphere transcriptomes alpha diversity and structural changes of bacteria and fungi under *G. yamadae* invasion. (a) The alpha diversity of bacterial and fungal transcriptomes in diseased leaves and healthy leaves across six developmental stages of crabapple rust disease. Box plots show the range of estimated values between the 25th and 75th percentiles, with the median, minimum, and maximum observed values within each dataset. Different letters indicate statistically significant differences was determined using one-way ANOVA with Tukey-HSD post hoc test (*P* < 0.05) or Kruskal-Wallis with Wilcoxon’s test (*P* < 0.05). (b) The correlation of Shannon index between bacterial and fungal transcriptomes. Solid lines represent the results of the simple linear regression models, while the surrounding grey band represents the 95% confidence interval. (c) PCoA of bacterial and fungal transcriptomes based on the Bray-Curtis distance matrix, R² and *P* were calculated using PERMANOVA test. Asterisks show the *P* significance level, \**P* < 0.05, \*\**P* < 0.01, \*\*\**P* < 0.001 and \*\*\*\**P* < 0.0001 and NS denotes no statistical significance.

We found that phyllosphere bacterial, fungal and viral transcriptome compositions were significantly influenced by three main factors: leaf condition (healthy or diseased), sampling stage, and their interaction. Among these, leaf condition emerged as the most significant determinant of transcriptome composition across all microbial groups (bacterial, fungal, and viral; Figure 1c, S1c; Table 1, S2b). To better understand the compositional differences, we employed Principal Coordinate Analysis (PCoA) based on Bray-Curtis distance. This analysis highlighted the distinct clustering of microbial transcriptomes across conditions and stages. For fungal and viral transcriptomes, the primary axis (PCoA 1) accounted for 46% and 48% of the total variance, respectively, delineating the overall differences in transcriptome composition. Separation based on leaf condition (healthy vs. diseased) was observed along the secondary axis (PCoA 2), which explained an additional 33% of the variation. Further analysis showed that bacterial transcriptomes in diseased leaves exhibited the highest Bray−Curtis dissimilarity during the early stage of infection (Stage 1). In contrast, viral communities displayed peak dissimilarity during the final stage of disease progression (Figure S1d).

**Table 1.**
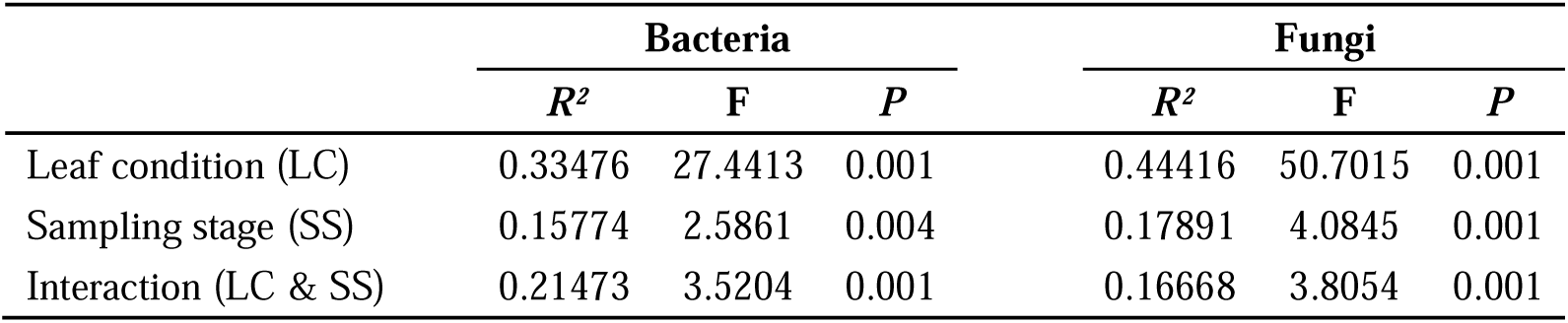
The effects of leaf condition and sampling stage on the structure of bacterial and fungal transcriptomes based on PERMANOVA with 999 permutations.

### 2.2 Overview of the composition and differential transcripts of phyllosphere transcriptomes

We characterized the composition and differential taxonomic profiles of phyllosphere transcriptomes in response to *G. yamadae* infection (Figure 2). The infection substantially altered the transcriptomic landscape of the crabapple phyllosphere. In healthy leaves, bacterial transcriptomes consistently dominated, typically accounting for 50% or more of the total transcriptomes across all stages (Figure 2a). Except for the first stage and the sixth stage, the relative expression abundance of microbial groups followed this order: bacteria > fungi > viruses. Conversely, in diseased leaves, fungal transcript abundance progressively increased, peaking at 96.2% in the final stage. Concurrently, the relative expression abundance of bacterial and viral transcriptomes decreased, although bacterial transcripts remained more abundant than viral ones throughout the progression. Additionally, archaeal transcripts were exclusively detected in the final diseased stage.

**Figure 2.**
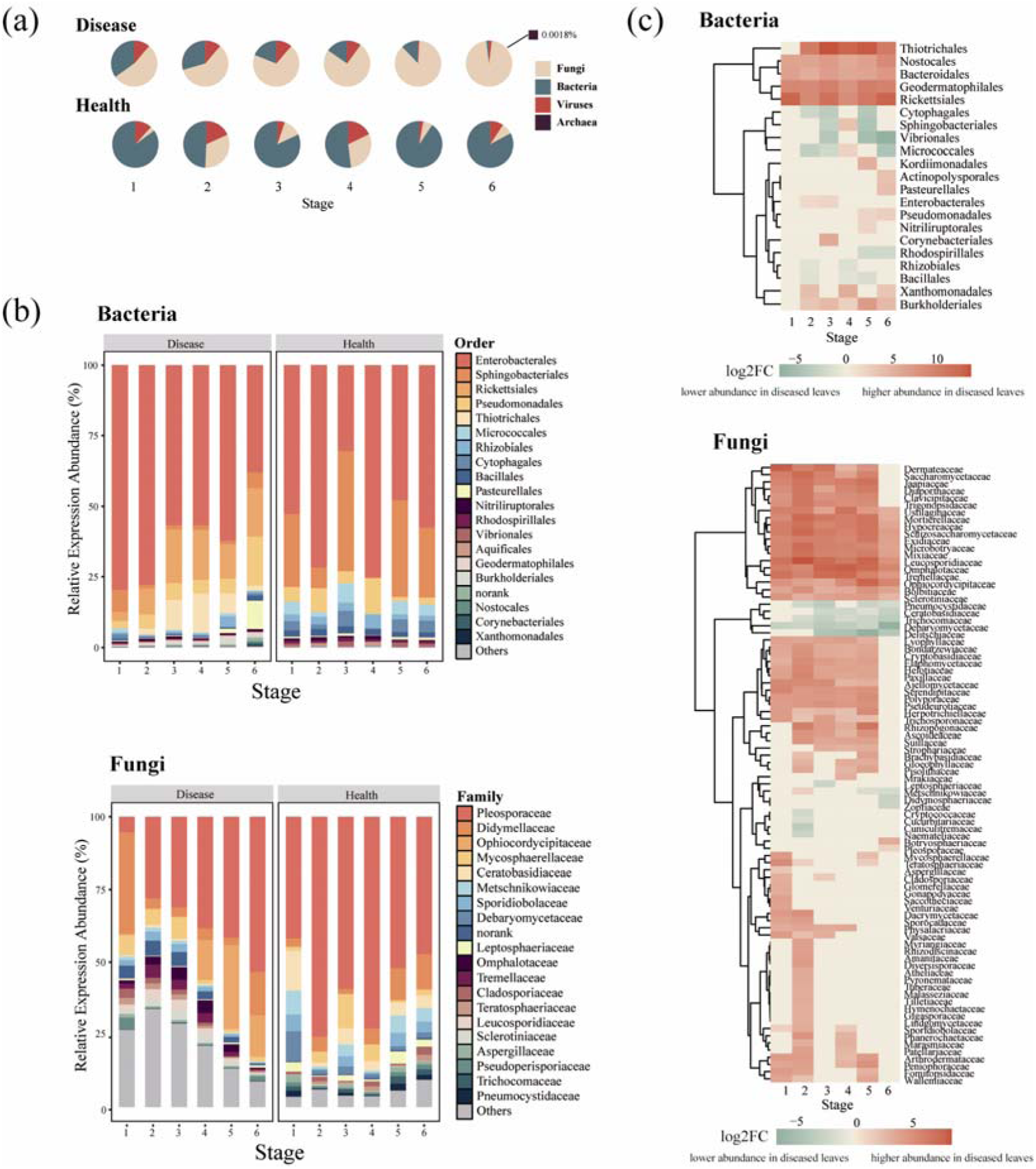
Overview of the composition and differential transcripts of phyllosphere transcriptomes. (a) The composition of phyllosphere transcriptomes in different stages, showing the relative abundance of bacterial, fungal, and viral transcriptomes. (b) Taxonomic compositions of the bacterial transcriptome at the order level and the fungal transcriptome at the family level, highlighting the predominant groups in both healthy and diseased leaves at different stages of infection. (c) Significantly differentially expressed transcripts in crabapple leaves under different conditions. Differential expression analysis was performed using a generalized linear model (GLM) to identity transcripts showing significant differences at each developmental stage (*P* < 0.05, FDR corrected).

Taxonomical classification of the bacterial transcriptomes at the order level and fungal transcriptomes at the family level reveals key shifts in microbial composition based on transcriptomic profiles, highlighting the top 20 taxa transcripts by relative expression abundance (Figure 2b). In diseased leaves, the dominant bacterial taxa included Enterobacterales (63.4%), Rickettsiales (12.1%), Pseudomonadales (6.6%), Thiotrichales (5.1%) and Sphingobacteriales (3.7%). In the healthy leaves, the most abundant bacterial orders were observed for Enterobacterales (51.5%), Sphingobacteriales (27.9%), Micrococcales (4.3%), Cytophagales (3.9%) and Pseudomonadales (3.9%). Several bacterial orders, including Thiotrichales, Nostocales, Bacteroidales, Geodermatophilales, Rickettsiales, Xanthomonadales and Burkholderiales, showed significant upregulation in diseased samples compared to healthy leaves (Figure 2c, S2). Conversely, Cytophagales, Vibrionales, Rhodospirillales, Rhizobiales and Bacillales, were significantly downregulated in diseased samples (*P* < 0.05).

For fungal transcriptomes, the family Pleosporaceae dominated both diseased and healthy leaves, accounting for 48.2% and 64.6%, respectively (Figure 2b). Other families with higher relative abundance in diseased samples included Ophiocordycipitaceae and Didymellaceae. Fungal families represented by Dermateaceae were notably upregulated in diseased leaves compared to healthy leaves and clustered together in the hierarchical analysis (Figure 2c). Unlike the bacterial communities, fungal transcripts generally showed increased expression during rust infection, with relatively fewer taxa downregulated. Interestingly, some bacterial taxa exhibited stage-specific differential expression patterns. Sphingobacteriales and Micrococcales were significantly upregulated in the third diseased stage and downregulated in the early and late stages. However, no similar stage-dependent trends were observed in fungal communities.

### 2.3 Changes in phyllosphere gene expression upon G. yamadae infection

To determine the gene expression response during each stage of *G. yamadae* infection, we quantified and visualized the number of expressed genes (Figure 3a). Overall, the total number of expressed genes, as well as stage-specific gene expression, was lowest at the initial stage of rust infection and peaked at the final stage. Pairwise comparisons of phyllosphere transcriptomes between diseased and healthy crabapple leaves were conducted at each stage to identify differentially expressed genes (DEGs). Transcripts with significant differential expression (log_2_Fold-Change ≥ 1 or ≤ −1, adjusted *P*-value < 0.05) were categorized DEGs (Figure 3b; Table 2, S3a). As the rust spots expanded, there was a marked increase in gene expression, with the number of upregulated genes consistently surpassing the number of downregulated genes at each stage.

**Figure 3.**
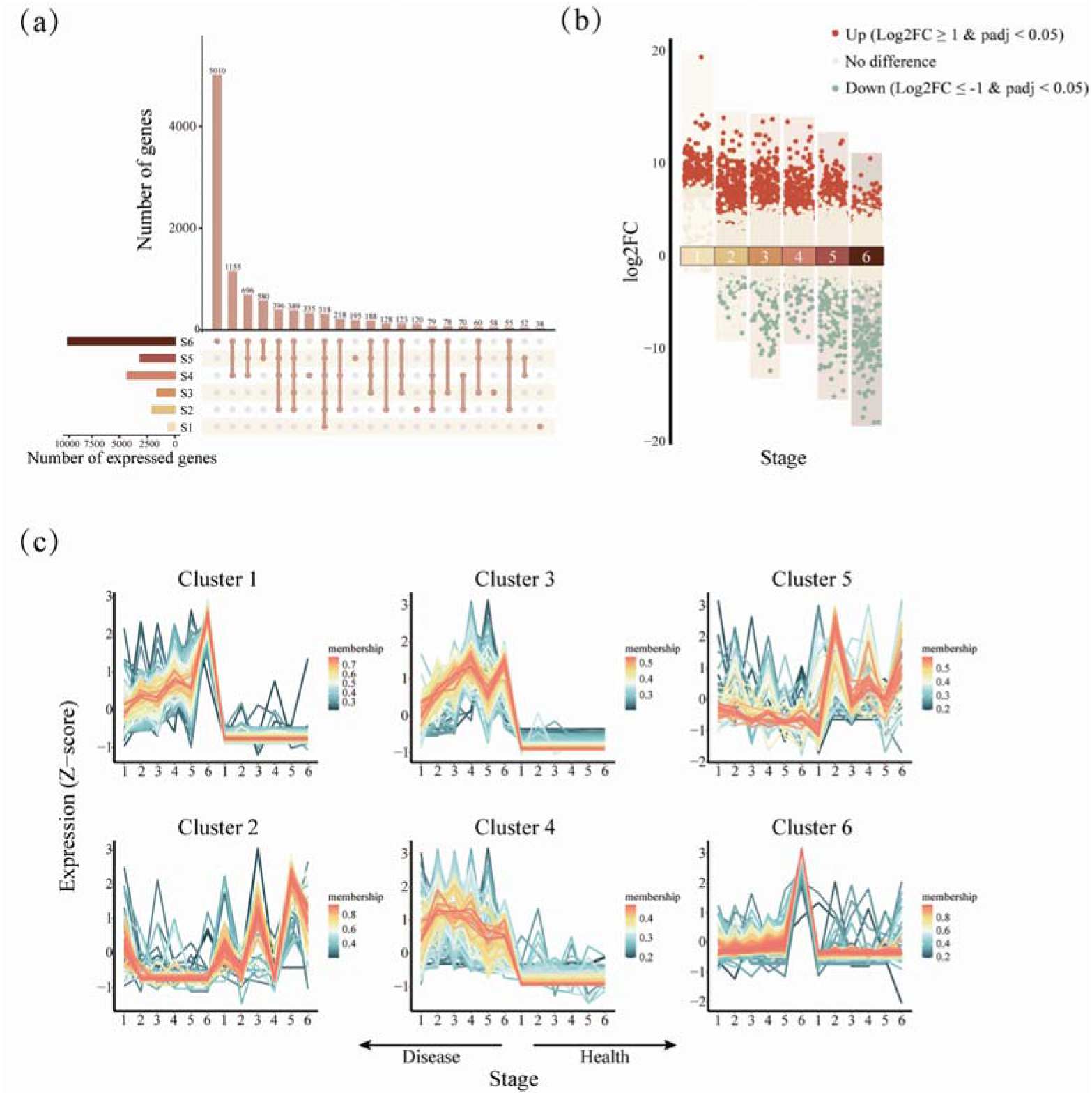
Expression profiling of phyllosphere target genes infected with *G. yamadae*. (a) The overlap of expressed genes across different stages of rust disease, showing top 22 sets. (b) Differentially expressed genes (DEGs) between diseased and healthy leaves at each stage of infection. Red dots indicate upregulated genes in diseased samples compared to healthy samples (log_2_Fold-Change ≥ 1, adjusted *P*-value < 0.05), green dots indicate downregulated genes in diseased samples compared to healthy samples (log_2_Fold-Change ≤ −1, adjusted *P*-value < 0.05), and the yellow dots denote genes with no significant difference between diseased and healthy samples. (c) Expression profile of DEGs clusters through c-means clustering. The color scale indicates the degree of membership of each gene to the respective clusters based on expression patterns, with darker colors representing higher membership to a given cluser.

**Table 2.**
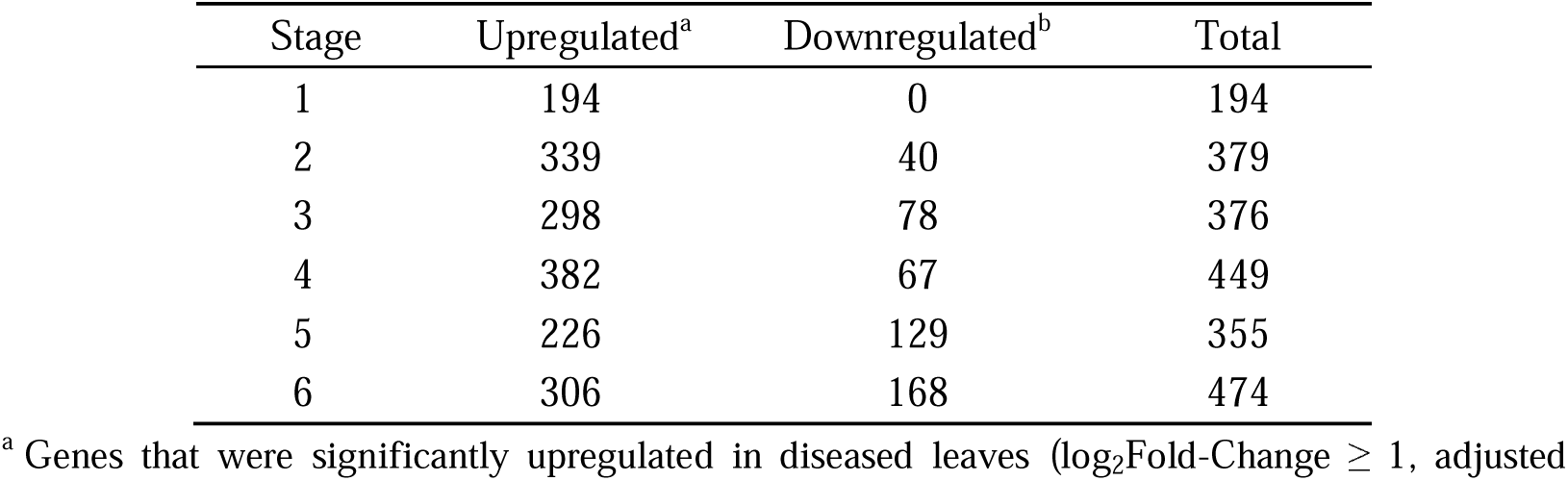
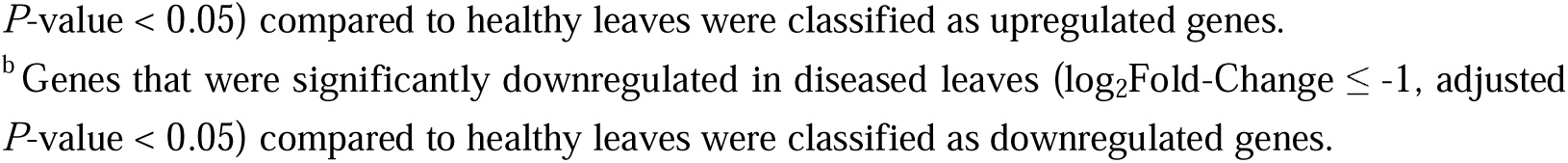
The results of differential gene expression analysis in crabapple phyllosphere at each stage.

| Stage | Upregulated<sup>a</sup> | Downregulated<sup>b</sup> | Total |
| --- | --- | --- | --- |
| 1 | 194 | 0 | 194 |
| 2 | 339 | 40 | 379 |
| 3 | 298 | 78 | 376 |
| 4 | 382 | 67 | 449 |
| 5 | 226 | 129 | 355 |
| 6 | 306 | 168 | 474 |

<sup>a</sup> Genes that were significantly upregulated in diseased leaves ( $\log_2\text{Fold-Change} \geq 1$ , adjusted
$P$ -value $< 0.05$ ) compared to healthy leaves were classified as upregulated genes. <sup>b</sup> Genes that were significantly downregulated in diseased leaves ( $\log_2\text{Fold-Change} \leq -1$ , adjusted $P$ -value $< 0.05$ ) compared to healthy leaves were classified as downregulated genes.

To further investigate the characteristics of the DEGs that actively respond to *G. yamadae* infection, we performed clustering analysis using the c-means clustering method and visualized the expression profiles of each cluster (Table S3b; Figure 3c). The six resulting clusters exhibited distinct expression patterns and taxonomic compositions. In particularly, Cluster 5 exhibited high expression in healthy leaf tissues. The *Klebsiella* genus was the predominant transcript in both diseased and healthy samples, accounting for 61% and 22.9% respectively, with all identified as *Klebsiella pneumoniae*. Additionally, *Sphingobacteriaceae bacterium* was a major contributor in healthy leaves, comprising 51% of the total transcripts. Clusters 1, 3 and 4 showed similar expression trends but differed significantly in their taxonomic composition. Cluster 1 contained genes with highest relative expression abundance in diseased leaves, primarily from viruses and bacteria, such as *Foveavirus* (33.3%), *Acinetobacter* (12.9%), and *Klebsiella* (11.3%). In contrast, Clusters 3 and 4 were predominantly composed of fungal transcripts. In cluster 3, *Monilinia* accounting for 10.2% of the relative expression abundance in diseased samples, whereas Cluster 4 was mainly composed of *Alternaria*, which represented 40.5% of the total transcripts.

### 2.4 Dynamic functional profiles of phyllosphere transcriptomes across different stages

To investigate the impact of rust infection and disease progression on the functional attributes and activities of the crabapple phyllosphere microbiota, we annotated the transcripts for functional analysis (Figure 4, S3, S4). Of the target unigenes retrieved from the sequence and bioinformatics data, 29.5%, 52.7% and 2.8% were assigned to the KEGG (Kyoto Encyclopedia of Genes and Genomes), GO (Gene Ontology) and CAZy (Carbohydrate-Active enZYmes) databases, respectively.

**Figure 4.**
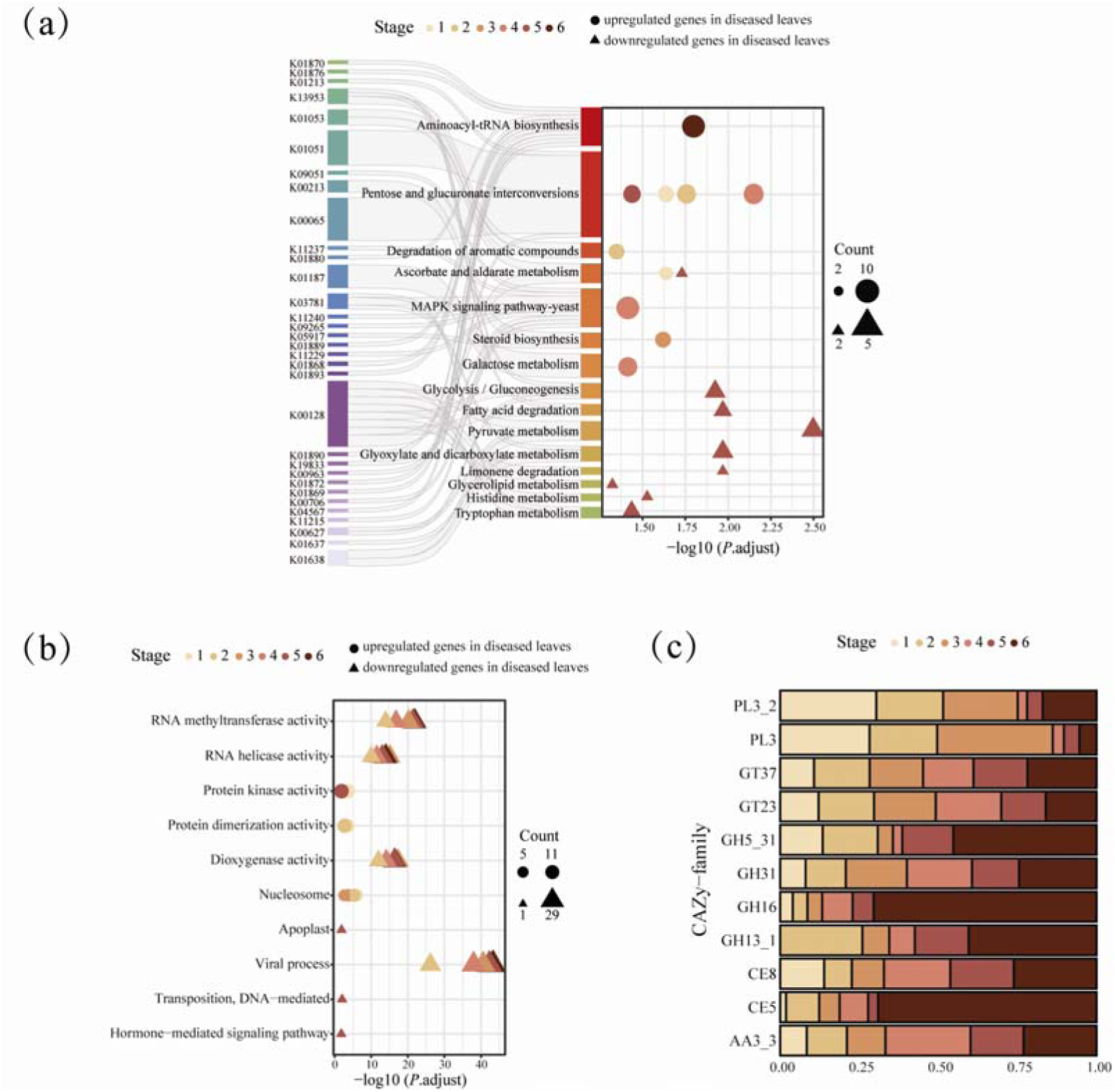
Functional annotation of phyllosphere transcripts in crabapple leaves based on the Kyoto Encyclopedia of Genes and Genomes (KEGG), Gene Ontology (GO) and Carbohydrate-Active enZYmes (CAZy) databases. (a) and (b), The KEGG pathway and GO functional module enrichment analysis of all DEGs based on the hypergeometric test, respectively. The *X*-axis represents the enrichment significance of the pathway or module (*P* < 0.05, FDR corrected, log2 transformed). The circle and triangle sizes represent the number of enriched genes. (c) The relative abundance of significantly differentially expressed carbohydrate-active enzymes (CAZy) across rust disease stages. Differential expression analysis based on generalized linear model (GLM) was used to identity CAZy families showing significant differences at each diseased stage (DESeq2; *P* < 0.05, FDR corrected).

After filtering out annotations unrelated to phyllosphere microorganisms, we performed KEGG and GO enrichment analysis on DEGs at each stage and identified significantly enriched pathways (Figure 4a). Overall, the active pathways in phyllosphere microorganisms of crabapple under infection by *G. yamadae* were primarily involved in primary metabolic processes, with almost all KEGG classifications increasing due to rust disease (Figure 4a, S3). Although the metabolism-related classifications did not exhibit overall significant temporal trends in diseased leaves, some specific metabolic pathways showed significant patterns compared to healthy samples. For example, the pentose and glucuronate interconversions pathway was enriched almost throughout the entire disease progression (Figure 4a). Conversely, transcripts associated with RNA methyltransferase activity, RNA helicase activity, dioxygenase activity and viral process were significantly downregulated during the complete course of rust disease (Figure 4b). Specifically, at the onset of the disease, certain transcripts related to carbohydrate metabolism were notably enriched, such as those involved in ascorbate and aldarate metabolism category (Figure 4a). Simultaneously, pathways involved in the degradation of aromatic compounds were active during the second stage of infection (Figure 4a). Transcripts associated with cellular components affecting gene expression and regulation, such as nucleosome, as well as those related to protein kinase and dimerization activity, were significantly upregulated in the early stages of rust disease (Figure 4b). As lesions expanded, pathways including steroid biosynthesis, galactose metabolism and MAPK signaling were significantly enriched, alongside sustained activity of nucleosome (Figure 4a, 4b). Interestingly, a few categories showed significant upregulation in the late stages of rust disease, such as the aminoacyl-tRNA biosynthesis pathway related to translation, and genes associated with protein kinase activity also exhibited significant enrichment in the latest stage of the disease progression. In addition to the pathways and molecular functions mentioned above, which were mostly downregulated across all diseased stages, certain functions were significantly downregulated only in the late stages of rust disease, including carbohydrate metabolism, lipid metabolism, amino acid metabolism, and apoplastic activity (Figure 4a). Notably, some pathways exhibited temporal patterns, with the ascorbate and aldarate metabolism pathway enriched in the first stage of *G. yamadae* infection but suppressed in the fifth stage.

To gain a more detailed resolution of specific functions associated with metabolism-related pathways, we searched for carbohydrate-active enzymes (CAZy) database. Similar to the KEGG analysis results, the diversity of KEGG orthology (KO) and CAZy families showed similar temporal patterns, and *G. yamadae* infection also led to an increase expression of all CAZy families (Figure S4a, S4b). After identifying for differentially expressed CAZymes across various disease stages, we focused on 11 CAZy families actively involved in regulation (Figure 4c, S4c). Remarkably, certain glycoside hydrolases, potentially related to fungal cell wall component degradation, exhibited increased activity in diseased leaves. These include enzymes involved in the degradation of chitin (GH5), glucans (GH5, GH16 and GH13) and mannans (GH5 and GH31). The relative expression abundance of these glycoside hydrolases (GH5, GH16, and GH13) increased as the lesions expanded and became more enriched in the late stages of rust disease (Figure 4c). Additionally, CAZy families that may interact with the host plant were also enriched in the infected leaves at different stages, including enzymes that participated in the degradation of plant cell wall components such as pectin (PL3 and CE8), cellulose (AA3), lignin (AA3), and xylan (CE5) (Figure 4c, S4c). Among these, the AA3 and CE5 families were upregulated in early and middle stages of rust disease, similar to the patterns of glycoside hydrolases. In contrast, PL3 and CE8 families were upregulated in the late stages (Figure S4c).

### 2.5 Dynamic changes in the complexity of phyllosphere microbiome functional co-occurrence networks as lesions expanded

To assess how lesion expansion impacts interactions among functional genes in the phyllosphere, we conducted co-occurrence network analysis and calculated topological properties for both healthy and diseased leaves at various stages (Figure 5). In all groups, functional genes related to microbial metabolism were the most predominant category (Figure 5a). The infection of *G. yamadae* altered the complexity of the phyllosphere co-occurrence networks, with network complexity indices showing opposite trends between healthy and diseased samples. Specifically, diseased leaves exhibited higher values in the number of nodes, number of edges, and average degree indices compared to healthy leaves, with these indices gradually increased as lesions expanded (Figure 5b). The modularization index, initially higher in diseased during the early stages of infection than in healthy samples, decreased progressively throughout the rust disease progression, eventually falling below the levels seen in healthy leaves by the middle and late stages of infection. In contrast, network density and clustering coefficient indices were higher in healthy leaves at early stages, but elevated in diseased samples as the infection advanced to the middle and late stages.

**Figure 5.**
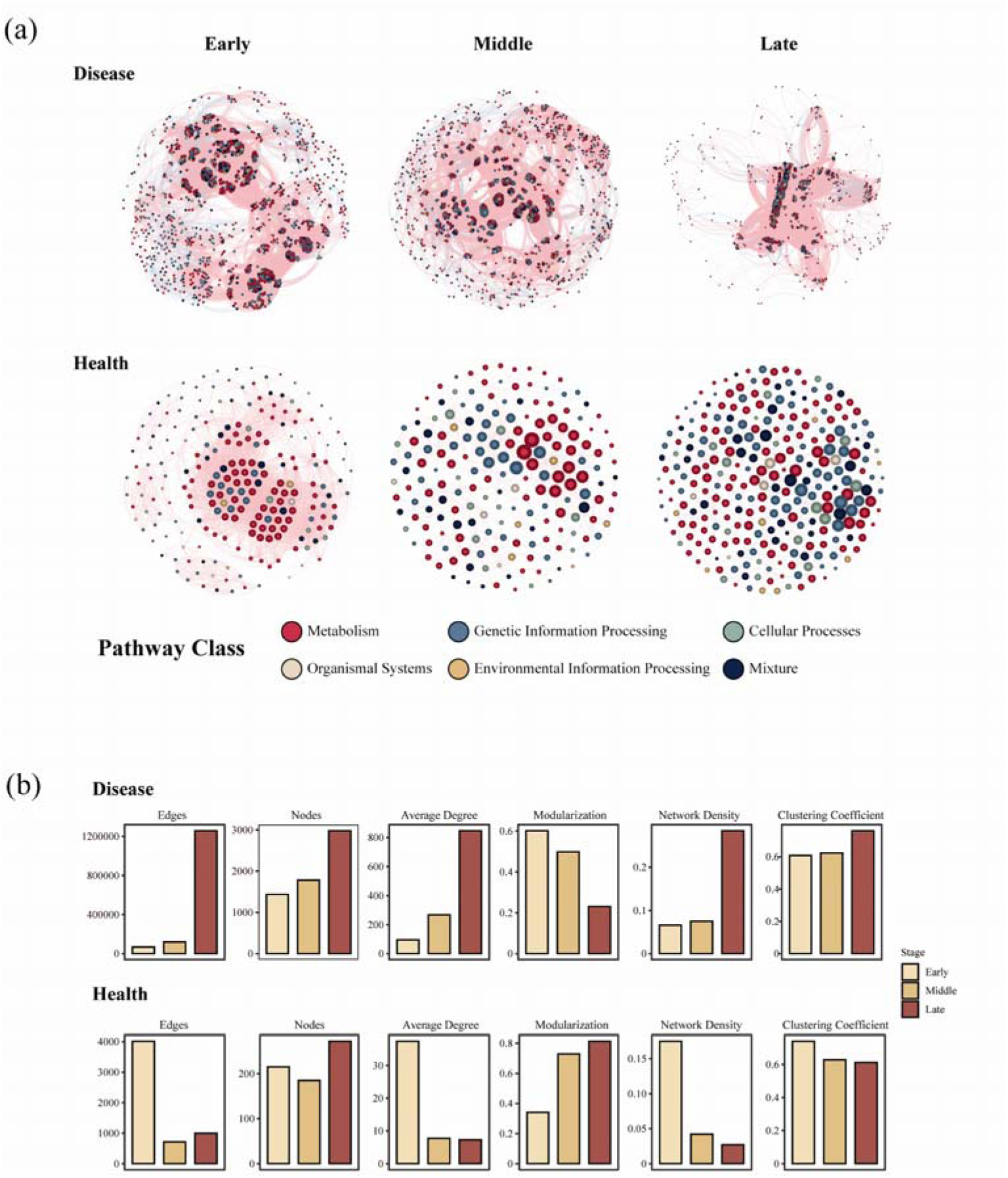
The co-occurrence networks of phyllosphere microbiome functional genes in diseased and healthy samples at different stages of rust disease. (a) Co-occurrence networks of functional genes in diseased and healthy samples at early, middle, and late stages of infection. The nodes in the networks are colored according to functional annotations derived from the KEGG database. Positive correlations between genes are represented by red edges, while negative correlations are indicated by blue edges. (b) Topological properties of the functional gene co-occurrence networks for both diseased and healthy samples across different stages of disease progression.

### 2.6 Contribution of phyllosphere functional genes to the pathogenesis of crabapple rust disease

To identify transcripts actively involved in the pathogenesis of crabapple rust disease, we conducted a random forest analysis, focusing on transcripts related to *G. yamadae* (order Pucciniales) and their contribution to disease progression (Figure 6; Table S4). Out of the 34 transcripts identified as significant important for predicting Pucciniales abundance (*P* < 0.05), *Alternaria alternata* emerged as the most predominant species. These key transcripts were primarily involved in several critical biological processes and pathways: the MAPK signaling pathway, aminoacyl-tRNA and steroid biosynthesis, as well as the production of CAZy families that target plant cell wall components. Specifically, enzymes that degrade pectin (PL3), xylan (CE5), and cellulose (AA3) were linked to this functional activity. Additionally, *Pseudovirgaria hyperparasitica* was found to contribute significantly to the production of pectin-degrading enzymes (CE8), which are involved in breaking down the plant cell wall. Transcripts involved in steroid biosynthesis process were primarily attributed to *Mortierella elongata*, and these transcripts were identified as the most important contributor to changes in Pucciniales abundance. Other metabolism-related transcripts also showed strong significance for Pucciniales dynamics. For instance, *Microbotryum intermedium* played a role in the degradation of aromatic compounds and ascorbate and aldarate metabolism, *Jaapia argillacea* was involved in galactose metabolism, and *Termitomyces* sp. J132 played a role in pentose and glucuronate interconversions. Interestingly, several transcripts associated with fungal cell wall degradation also showed significant importance in the model, including genes involved in the degradation of glucans (e.g., from *Saitozyma podzolica* and *Alternaria tenuissima*) and mannans (e.g., from *Alternaria tenuissima* and *Jaapia argillacea*). These findings underscore the significance of microbial interactions and enzymatic activities in contributing to the pathogenesis and progression of crabapple rust disease.

**Figure 6.**
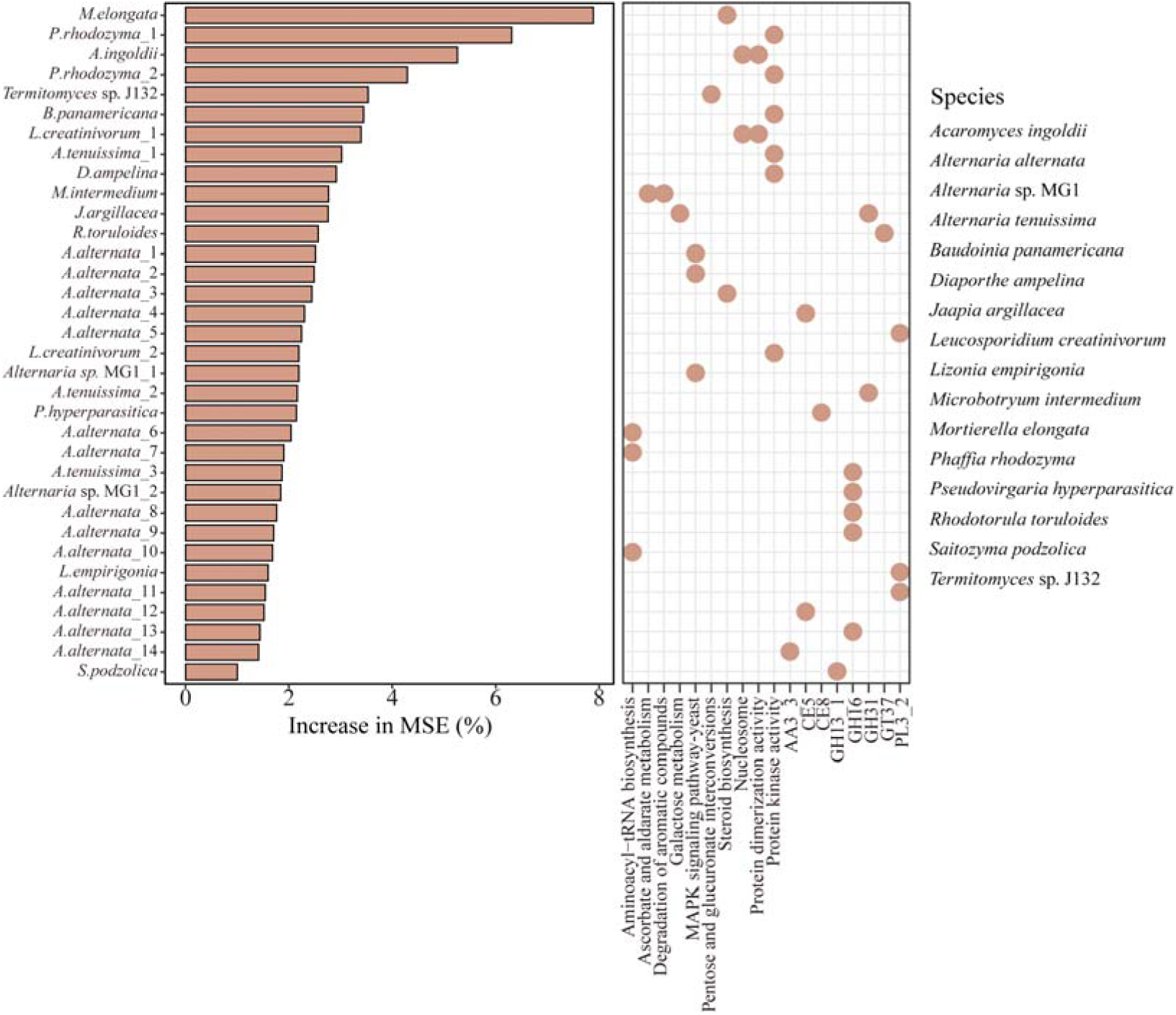
Importance ranking of phyllosphere microbial functional genes in the severity of crabapple rust disease determined by random forest analysis. The bar plot on the left shows the importance ranking of functional genes, with the *X*-axis representing the percentage increase of mean square error (MSE), and the y-axis represents functional gene identifiers. The higher the percentage increase in MSE, the more important the gene is in predicting pathogen severity. The dot plot on the right displays the specific functions associated with these genes, where the x-axis represents the functional categories and the y-axis lists the gene identifiers. All functional genes shown in this plot were statistically significant in the random forest model (*P* < 0.05).

## 3 Discussion

### 3.1 Rust disease shapes phyllosphere microbiome assembly in crabapple

The assembly and composition of the plant microbiome are intricately linked to the plant health, with pathogen infections often resulting in dramatic shifts in microbial composition and functional strategies across different plant-pathogen systems, potentially influencing the host resistance or susceptibility to disease ^[3]^. In our study, we observed a notable increase in species richness across bacteria, fungi, and viruses, which occurred following *G. yamadae* infection in crabapple leaves, coupled with more active microbial expression patterns in diseased leaves (Figure 1a, S1a, S1b). These shifts could reflect a complex interplay, where the plant may recruit antagonistic microbes to counter the pathogen or, alternatively, pathogen-associated microbes could be facilitating disease progression by occupying ecological niches vacated by stressed or damaged host tissues ^[8]^.

An intriguing observation in our study was the negative correlation between bacterial and fungal diversity in response to pathogen infection (Figure 1a,1b), suggesting competitive interactions between these two major microbial groups, potentially driven by pathogen-induced shifts in the phyllosphere microbiome. As *G. yamadae* colonizes the leaf surface and expands lesions, this could create selective pressures that favor the proliferation of certain fungal species, potentially outcompeting the bacterial populations. Such a competitive dynamic would be consistent with the ecological imbalance induced by the rust pathogen, where opportunistic fungi associated with the pathogen gain a competitive edge. Further support for this competitive dynamic is provided by changes in community evenness: as lesions expanded, bacterial species displayed more uniform expression abundance (increased evenness), while the fungal species showed greater divergence in expression abundance (decreased evenness) (Figure S1b). We postulate that this divergence in fungal community evenness may be induced by pathogen invasion, which disrupts the host’s defense mechanisms. This allows certain epiphytic fungi, particularly those associated with the pathogen, to rapidly colonize and infect the compromised host tissues, thereby putting bacterial communities at a competitive disadvantage, especially in terms of nutrient acquisition ^[8]^. This hypothesis is reinforced by the observed patterns in the relative expression abundance across the microbial groups in the phyllosphere: bacteria consistently dominated across all stages in healthy leaves, whereas fungi progressively emerged as the predominant microbial group in diseased leaves, with their relative abundance peaking as the lesions expanded (Figure 2a). These findings underscore the increasing importance of fungi—both opportunistic pathogens and symbiotic microbes—in the pathogenesis process. As fungal populations gain prominence in diseased tissues, they may not only exacerbate the disease but also shift the overall microbial landscape, influencing the host’s response to infection and potentially altering disease outcomes.

### 3.2 Shifts in microbial community composition and functional groups

PCoA results revealed that microbial community composition in the phyllosphere was strongly influenced by leaf condition, with distinct microbial signatures found in diseased leaves versus healthy leaves (Figure 1c, S1c, S1d). Interestingly, during the early and middle stages of infection, bacterial diversity was lower than in healthy leaves at corresponding stages, however, in the late stages of infection, bacterial diversity in the diseased leaves surpassed that in healthy leaves (Figure 1a). This suggests an adaptive restructuring of the microbial community over the course of the infection, where specific taxa may have been displaced or outcompeted initially, but later stages of infection provide new ecological niches, allowing for the gradual re-establishment of bacterial communities. This restructuring may reflect the shifting availability of nutrients or other ecological factors that favor certain bacterial taxa over time ^[18]^.

The infection of *G. yamadae* significantly altered the composition of the phyllosphere microbial community, and the profiles of relative expression abundances revealed notable shifts (Figure 2). While the dominant bacterial taxa at the order level were relatively similar in both healthy and diseased samples, certain transcripts with low cumulative relative abundance exhibited significant upregulation or downregulation in diseased leaves (Figure 2b, 2c, S2). This suggests that pathogen infection triggers a distinct microbial response in terms of gene expression, likely reflecting shifts in microbial activity associated with disease progression. In the fungal community, the family *Pleosporaceae* was the dominant group in both disease and healthy leaves, with most fungal transcripts generally upregulated under pathogen infection, particularly when compared to bacterial community. This suggests a greater involvement of fungi in the host response to pathogen infection, possibly through interactions that exacerbate or modulate pathogen development ^[42]^.

We identified several microbes in both the bacterial and fungal communities with potential antagonistic characteristics (Figure 2, 3). For instance, certain members of the order Nostocales, such as *Nostoc calcicola* and *Nostoc linckia*, have been documented as antagonists capable of inhibiting *Fusarium oxysporum*-induced wilt in tomatoes ^[43]^. Additionally, strains of *Klebsiella* from the order Enterobacterales have been shown to suppress rust lesions in coffee leaves ^[44]^. The genus *Methylobacterium*, enriched in the healthy phyllosphere, has been associated with improved plant growth and reduced disease incidence ^[45]^. These antagonistic microbes, by limiting the growth or activity of pathogens, might help mediate plant defense responses and support overall plant health during pathogen exposure. Moreover, we observed groups with potential antagonistic effects within the fungal community. For example, members of *Mortierellaceae* family can promote plant growth and seed production of *Arabidopsis thaliana* by mediating the upregulation of plant’s hormone production and activating its defense responses against pathogens ^[46]^. In the present study, the transcripts of these microbes were significantly enriched in diseased samples, suggesting their potential role in enhancing the host’s ability to resist *G. yamadae* infection, where beneficial fungi or bacteria suppress pathogen proliferation and bolster host defenses ^[8]^.

In addition to these beneficial microbes, we also observed that fungal communities that gradually dominated the phyllosphere with lesion enlargement consisted primarily of well-known opportunistic pathogens. The upregulated taxa in this category included members of the families *Diaporthaceae*, *Clavicipitaceae,* and *Mycosphaerellaceae* ^[47–49]^. This suggests that these fungal groups may be opportunistically colonizing the highly vulnerable tissues of diseased leaves, exploiting the conditions created by *G. yamadae* infection to further promote disease spread. The opportunistic nature of these pathogens underscores the complexity of plant-microbe interactions in the context of disease, where both beneficial and harmful microbes dynamically interact and influence disease outcomes ^[8]^.

### 3.3 Functional attributes and microbial network complexity

Functional analysis of the phyllosphere microbiome revealed *G. yamadae* infection significantly influenced the functional attributes and activities of microbial communities, with a noticeable shift in the complexity of functional co-occurrence networks compared to those of healthy leaves. Specifically, the functional co-occurrence networks in *G. yamadae*-infected leaves exhibited more complex patterns, aligning with the increased network complexity observed in our prior co-occurrence network analyses at the taxonomic level (Figure 5) ^[8]^. This shift suggests an adaptive response by the microbial community to pathogen invasion, characterized by the reorganization of microbial interactions. Previous studies have highlighted that pathogen infection often leads to the formation of more intricate microbial networks a part of a defense strategy against the invading pathogen^[50,51]^. For example, more complex microbial networks have been reported in the phyllosphere of *Diaporthe citri*-infected citrus plants and in both the aboveground and belowground compartments of chili pepper affected by *Fusarium* wilt disease, compared to their healthy counterparts ^[20,30]^.These findings support the notion that increased network complexity may be a general response mechanism of the phyllosphere microbiome to external stress, aimed at maintaining community stability and enhancing functional redundancy under pathogen pressure. In our study, the increased functional network complexity in *G. yamadae*-infected samples likely reflects a coordinated response by the phyllosphere microbiome to counteract the pathogen’s effects. We hypothesize that this complexity reflects enhanced interactions among microbial functions that could be protective or antagonistic against the pathogen. Key topological metrics, such as modularity—a measure that often reflects the degree of specialized, compartmentalized interactions within a microbial community-further illustrate this dynamic shift. During the early stages of *G. yamadae* infection, we observed higher modularity in the functional networks of infected leaves compared to healthy ones (Figure 5b), which suggests that the microbial gene functions in diseased leaves were more stably correlated, potentially reflecting positive co-regulation with distinct microbial subgroups. These compartmentalized responses may be part of a plant defense, where specialized groups of microbes act in concert to reinforce the plant’s defense mechanisms against pathogen invasion. However, as the disease progressed, this stable co-regulation is disrupted, becoming less pronounced compared to healthy leaves, which evolved more stable functional networks over time ^[51]^.

### 3.4 Metabolic adaptations and enzymatic activities

The functional enrichment analysis provided critical insights into the mechanisms underlying the observed shifts in the phyllosphere functional co-occurrence networks during *G. yamadae* infection. For instance, DEGs were substantially enriched in carbohydrate metabolism pathways, including pentose and glucuronate interconversions pathway, ascorbate and aldarate metabolism, and galactose metabolism (Figure 4a). These findings suggest an upregulation of energy metabolism in the phyllosphere microbiome in response to the stress induced by *G. yamadae.* This increased activity in carbohydrate and energy-related pathways may be essential for microbial survival and proliferation during pathogen attack, potentially supporting the microbiome’s overall resilience under stressful conditions. Additionally, the pathogen also modulated microbial gene expression patterns, with nucleosome activity and dimerization activity being prominent in the early diseased stages, followed by the upregulation of the aminoacyl-tRNA biosynthesis pathway in later stages (Figure 4a, 4b). This dynamic regulation highlights the microbiome functional adaptation throughout the rust disease progression, likely aimed at enhancing survival and mitigating the adverse impact of pathogen ^[35]^.

Another key finding was the upregulation of degradation pathways involved in breaking down plant-produced antimicrobial substances. Specifically, we observed significant enrichment of genes associated with the degradation of aromatic compounds, such as those in the ascorbate and aldarate metabolism pathways (Figure 4a). This is consistent with our previous study on metabolic profiles of the host plants used in the same experiment setup, further emphasizing the complex interactions within the plant-pathogen-microbiome model ^[8]^. Such interactions are integral to the plant’s response to pathogen attack, as plants often recruit beneficial microorganisms with antagonistic functions against pathogens. *Rhizoctonia solani*-infected beet roots are enriched with *Chitinophagaceae* and *Flavobacteriaceae*, which secrete fungal cell wall-degrading enzymes to inhibit pathogen infection ^[21]^. Similarly, in our study, the secretion of fungal cell wall degrading enzymes (GH5, GH16, GH13, and GH31) that target cell wall components (chitin, glucan, and mannan) during the early and middle stages of rust disease supports this viewpoint (Figure 4c, S4c). These enzymes target key fungal cell wall components such as chitin, glucan, and mannan, providing a defensive mechanism against pathogen by directly undermining the structural integrity of the invading fungi. These findings underscore the critical role of microbial enzymatic activities in shaping plant-microbe interactions during *G. yamadae* infection. The secretion of fungal cell wall-degrading enzymes and the overall metabolic reprogramming of the microbiome may be vital strategies by which the phyllosphere microbiota responds to infection, either through direct antagonistic interactions with the pathogen or by supporting the host’s immune responses.

### 3.5 Role of beneficial microbes and pathobiome in disease dynamics

Random forest model further revealed important microbial taxa and their associated functional activities that significantly contribute to the regulation of microbial functions and pathogen dynamics during *G. yamadae* infection. Notably, we identified key microbes like *Jaapia argillacea* and *Saitozyma podzolica*, whose transcripts were essential in shaping microbial responses and modulating *G. yamadae* abundance (Figure 6). *S. podzolica*, a yeast known for its plant growth-promoting (PGP) abilities, has been shown to combat pathogens like *Fusarium oxysporum* f. sp. *Melongenae* through the secretion of fungal cell wall-degrading enzymes and antifungal metabolites ^[52]^. Additionally, we observed the enrichment of transcripts from beneficial microorganisms like *Mortierella elongata*, known for its involvement in steroid biosynthesis and its influence on *G. yamadae* abundance (Figure 6). *M. elongata* has been frequently identified as a PGP microorganism in soil and the rhizosphere, enhancing hormone production (e.g., IAA) and bolstering plant resistance to pathogens ^[53,54]^. These recruited beneficial microbes likely play dual roles, both in promoting plant health and in directly inhibiting pathogen proliferation.

In contrast to the beneficial microbes, we also detected the presence of certain pathogenic partners, or ‘pathobiomes’, that likely assist *G. yamadae* in facilitating the progression of disease, highlighting the complexity of the plant-pathogen-microbiome interplay ^[23]^. These microorganisms, typically part of the resident microbiota in healthy leaves, can be exploited by pathogens during infection. This phenomenon highlights how pathogens can “hijack” commensal or symbiotic microorganisms, converting them into allies that aid in overcoming host defenses and establishing infections. One key mechanism involves symbiotic microorganisms capable of producing plant cell wall-degrading enzymes, which can enhance pathogen colonization by breaking down structural components of the host plant tissues. Such activities have been observed in the infections of tomato and tobacco plants, where these enzymes facilitate pathogen entry and spread ^[25,26]^. Our analysis of CAZy indicated a significant enrichment of enzymes like pectate lyase (PL3), carbohydrate esterase (CE8), and acetyl xylan esterase (CE5) (Figure 4c). These enzymes are involved in the degradation of pectin and xylan, key polysaccharides that form part of the plant cell wall. Their upregulation throughout various stages of *G. yamadae* infection suggests that the pathogen and its associated microbiome are actively degrading plant cell walls, weakening host tissues and facilitating deeper colonization.

Despite functional redundancy observed across different microbial taxa, *Alternaria alternata* was identified as a primary contributor to the functional enrichment of microbial activities ^[55,56]^. Remarkably, *A. alternata* also participated in the MAPK signaling pathway, which is crucial for regulating spore formation, resistance to oxidative and osmotic stress, as well as for its pathogenesis in citrus ^[57]^. The results of GO functional module enrichment analysis further revealed active dynamics of phyllosphere microorganisms during the late stages of disease in the plant apoplast (Figure 4b). The plant apoplast, a key physical barrier against pathogens, play a significant role in triggering defense responses ^[28]^. Thus, we speculate that the dynamic shifts observed in microbial activities during late-stage infection may reflect increased opportunism by certain microbes, exploiting the deteriorating phyllosphere environment. However, the roles of these microorganisms in contributing to or mitigating disease progression require further experimental validation to elucidate their precise functions and interactions within this complex microbial ecosystem.

In summary, our study underscores the critical role of the phyllosphere microbiome in mediating plant-pathogen interactions and shaping disease outcomes. These insights provide a foundation for developing microbiome-based strategies to enhance plant health and resilience against pathogens.

## 4 Materials and Methods

### 4.1 Sample collection

Crabapple (*Malus* ‘Kelsey’) leaves were collected from trees located at the south gate of Olympic Park, Beijing (40°N latitude, 116.38°E longtitude) between June and September 2021. The sample process targeted both healthy (non-infected) and diseased (*Gymnosporangium yamadae*-infected) leaves from the same trees across six distinct stages of rust disease progression, ranging from the formation of spermogonia to the maturation of aecia. To ensure uniformity and control for genetic variation, three biological replicates were collected for each condition and stage. The leaf sample were immediately transported to the laboratory on dry ice immediately and stored at −80D until RNA extraction.

### 4.2 RNA extraction and metatranscriptomic sequencing

To prepare for RNA extraction, the leaf tissues were ground into a fine powder using RNase-free mortars and pestles with liquid nitrogen. Total RNA was extracted from the phyllosphere using the Fecal RNA Extraction Kit (Majorbio, Shanghai, China), following the manufacturer’s procedure. The integrity and concentration of the extracted RNA were measured with NanoDrop 2000 spectrophotometer (Thermo Scientific, MA, USA) and an Agilent 5300 Bioanalyzer (Agilent Technologies, Palo Alto, CA, USA). Ribosomal RNA (rRNA) was depleted using the RiboCop rRNA Depletion Kit for Mixed Bacterial Samples (Lexogen, USA), focusing on preserving mRNA for library preparation. For sequencing, 200 ng for each sample’s RNA was used for library preparation with the Illumina® Stranded mRNA Prep, Ligation (Illumina, San Diego, CA, USA). Paired-end sequencing was carried out on the Illumina Novaseq 6000 at Majorbio Bio-Pharm Technology Co., Ltd. (Shanghai, China).

### 4.3 Bioinformatics and statistical analysis

#### 4.3.1 Data preprocessing and quality control

Raw sequencing data were processed to ensure high quality and minimize contamination before further analysis. The sequencing data were processed using fastp v0.19.6^[58]^ for quality control, trimming adapter sequences from both 3’ and 5’ ends of reads. Readers shorter than 50 bp or those with an average base quality score below 20 were discarded to ensure high-quality data, and reads containing ambiguous bases (denoted as ‘N’) were also removed. Potential contaminant reads aligned to the host plant genomes were removed, including *Malus domestica* (GCF_002114115.1), *Malus baccata* (GCA_006547085.1), *Malus sylvestris* (GCF_916048215.2), and *Malus sieversii* (GCA_020795835.1) using BWA ^[59]^. To further eliminate non-target sequences, rRNA contamination was filtered using SortMeRNA ^[60]^, a tool designed to remove ribosomal RNA sequences. After cleaning the data, de novo transcript assembly was performed using Trinity v2.2.0 ^[61]^, a widely used tool for assembling RNA-seq data into longer transcript sequences without requiring a reference genome. Transcripts with lengths ≥ 300base pairs were retained for further analysis. The assembled transcripts were then de-replicated using CD-HIT v4.6.1 ^[62]^, setting an identity threshold of 0.95 and a minimum coverage of 0.9, which reduced redundancy and ensured that the final set of transcripts represented unique functional genes. The longest sequence was used as representative unigenes for downstream analysis.

#### 4.3.2 Taxonomic annotation

The assembled unigenes were aligned against the NCBI NR (non-redundant) database using DIAMOND v0.8.35 ^[63]^ with an e-value cutoff of 1e-5 to ensure reliable matches. Sequences identified as belong to Viridiplantae (plants), Metazoan (animals) and the pathogen (Pucciniales) were excluded from the analysis, allowing the focus to remain on the targeted microbial communities (Table S5). Transcript abundance was estimated using the RSEM v1.3.2 ^[64]^, generating FPKM (fragments per kilobase of transcript per million mapped reads) values for each sample, ensuring accurate comparisons of microbial activity in the phyllosphere microbiome.

#### 4.3.3 Diversity analysis

Alpha diversity indices, including Richness, Shannon and Pielou’s evenness, as well as beta diversity metrics such as Bray-Curtis dissimilarity, were calculated using the vegan package ^[65]^ in R. For statistical comparisons of alpha diversity indices (bacterial, fungal, and viral transcriptomes) between diseased and healthy leaves, one-way ANOVA with Tukey-HSD post hoc test or Kruskal-Wallis with Wilcoxon’s test was employed using the multcomp package ^[66]^. The Shannon index values for bacterial and fungal transcriptomes were correlated using linear regression models (Generalized Linear Models, GLM), implemented with the ggpmisc package ^[67]^. Beta diversity was assessed using Bray-Curtis distance matrices for different stages and leaf infection conditions, which were visualized through principal coordinates analysis (PCoA). Permutational multivariate analysis of variance (PERMANOVA) was applied to test the significant differences in community composition between sample groups using the vegan package ^[65]^.

#### 4.3.4 Differential expression and functional analysis

The expression matrix of the unigenes was filtered to retain genes expressed in all the consistently expressed across three biological replicates for each treatment. Genes that were expressed at each stage of rust disease was counted, and the intersection of these genes were visualized using the UpSetR package ^[68]^. Differentially expressed genes (DEGs), transcripts and carbohydrate-active enzymes (CAZymes) were identified using the DESeq2 package ^[69]^. Pairwise comparisons were made between adjacent sampling stages, applying a generalized linear model (GLM) approach with a significance threshold of *P* < 0.05 after false discovery rate [FDR] correction. Expression data, transformed to Z-scores and log-transformed FPKM values, were calculated using the c-means clustering algorithm in the TCseq package ^[70]^. For functional annotation, DIAMOND ^[63]^ was used to annotate with the KEGG and GO databases, and hmmscan v3.3.2 ^[71]^ was used for the CAZy database, with the alignment parameter e-value set to 1e-5 for both. The dynamic abundance patterns of KEGG classifications and CAZy families were presents based on normalized functional geng FPKM values and analyzed using one-way ANOVA test. Functional enrichment analysis of DEGs was performed using the clusterProfiler package ^[72]^. The unigenes obtained from KEGG and GO annotations were used as the background gene set to identify significantly upregulated and downregulated functions associated with DEGs at each time point, with FDR correction applied using the Benjamini-Hochberg method. The Spearman correlation of functional genes was calculated using the Hmisc package ^[73]^, and topological properties and the functional co-occurrence network were analyzed and visualized using Gephi ^[74]^ as disease lesions expanded. To identify functional genes most predictive of pathogen abundance, a random forest model we constructed used the rfPermute package ^[75]^. The relative abundance of Pucciniales was set as the response variable, with the expression levels of other functional gens serving as predictor variables. This model was employed to pinpoint microbial functional activities potentially linked to the pathogen’s colonization success.

## Acknowledgments

This study was supported by the National Natural Science Foundation of China (grant 32101527).

## Author Contributions

Q.X. conducted the experiments, analyzed the data, and edited the manuscript. Y.Z. conducted the experiments and edited the manuscript. S.T. designed the experiments, provided the experimental conditions, and contributed to the editing and review of the manuscript. All authors have read and approved the final version of the manuscript for publication.

## Data Availability Statement

The sequencing raw data have been uploaded to the National Genomics Data Center.

---

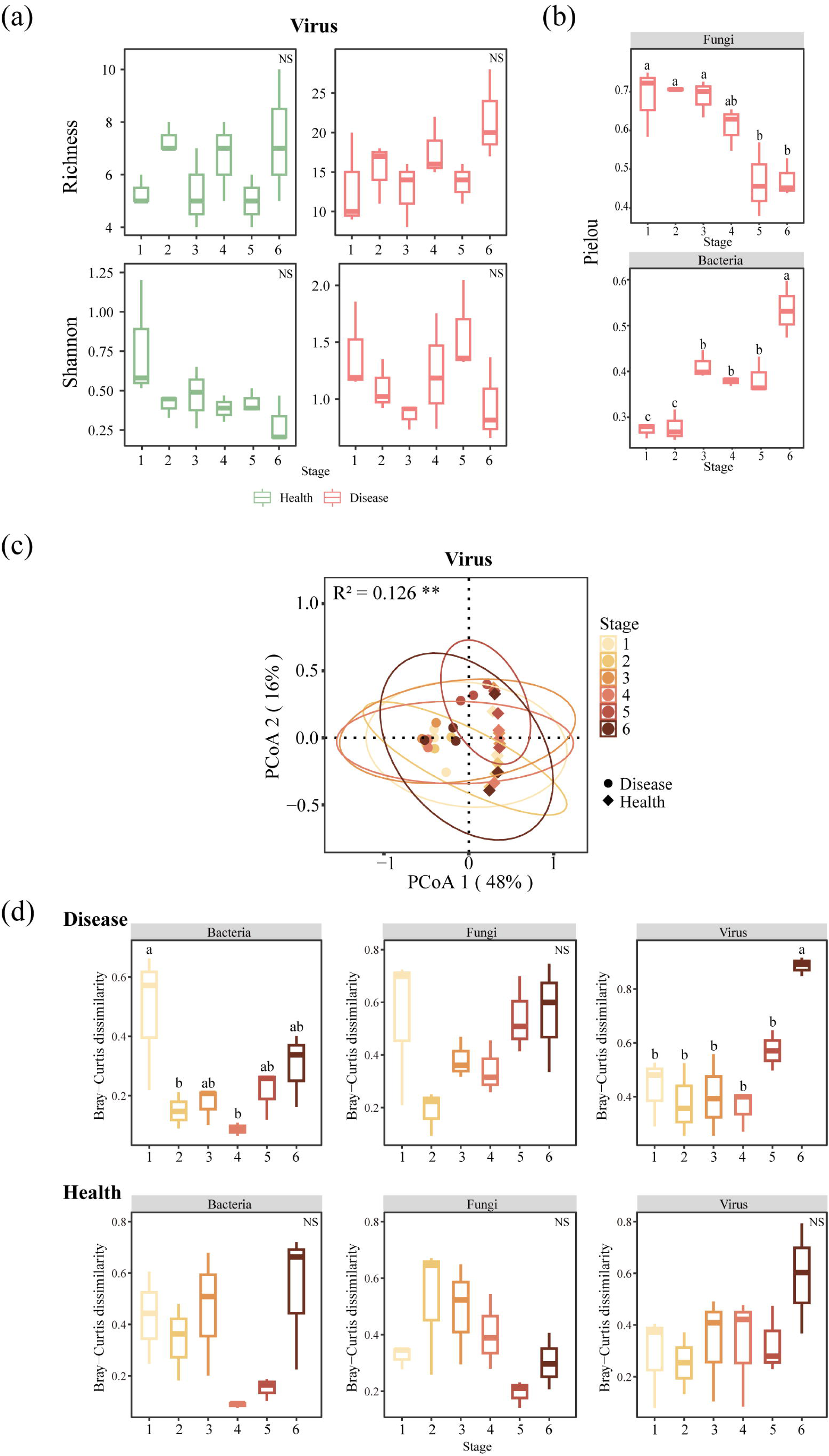

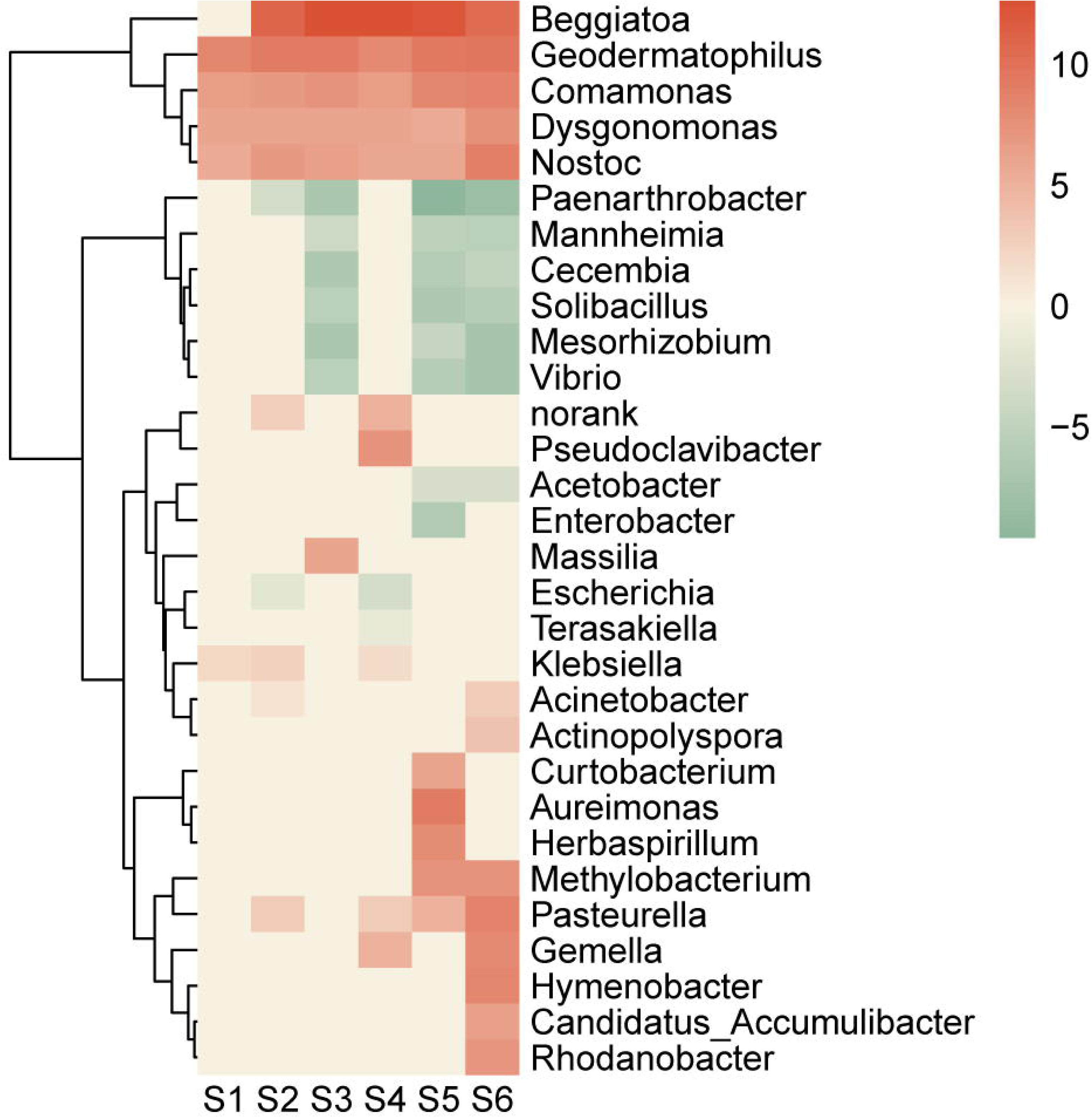

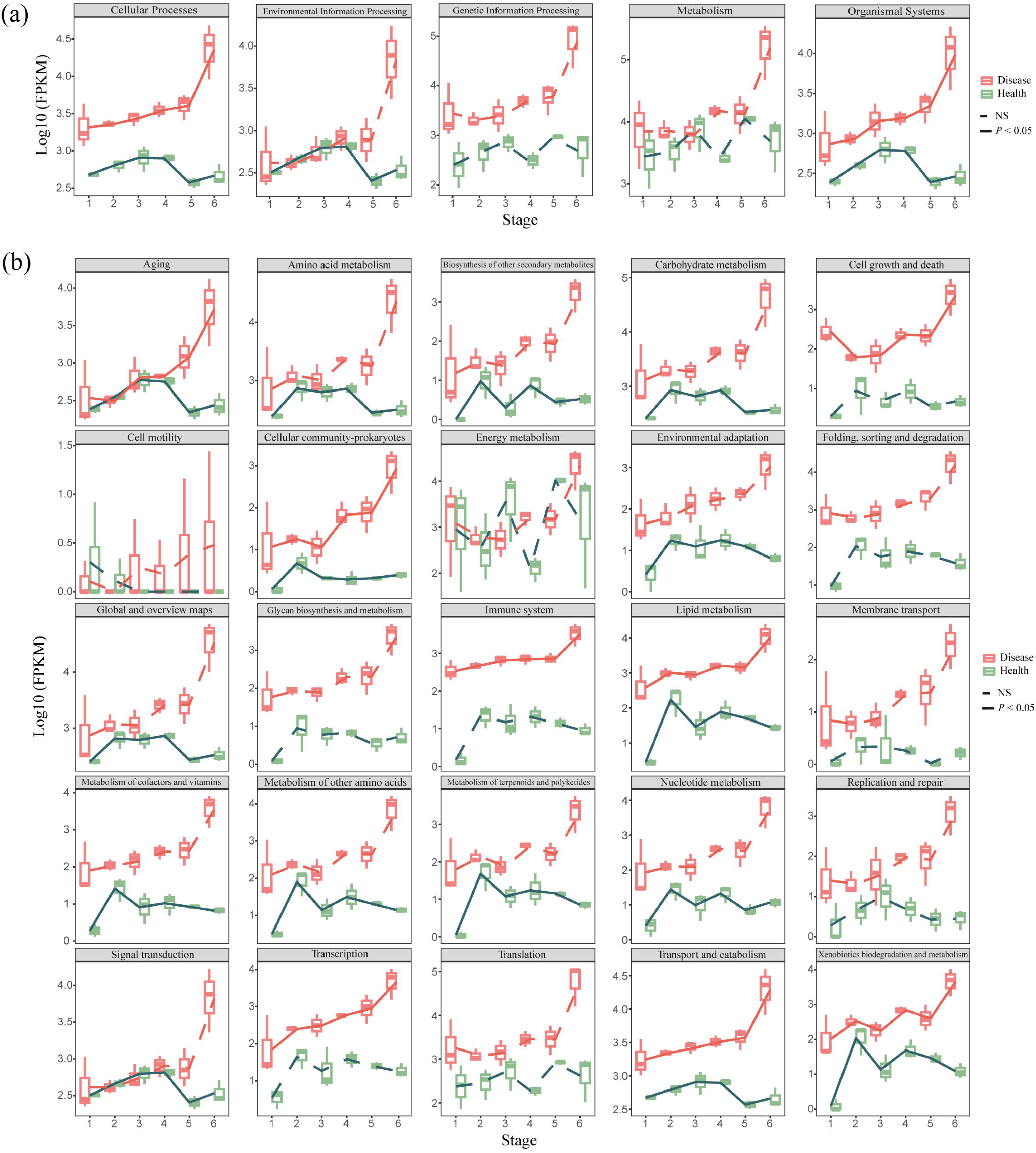

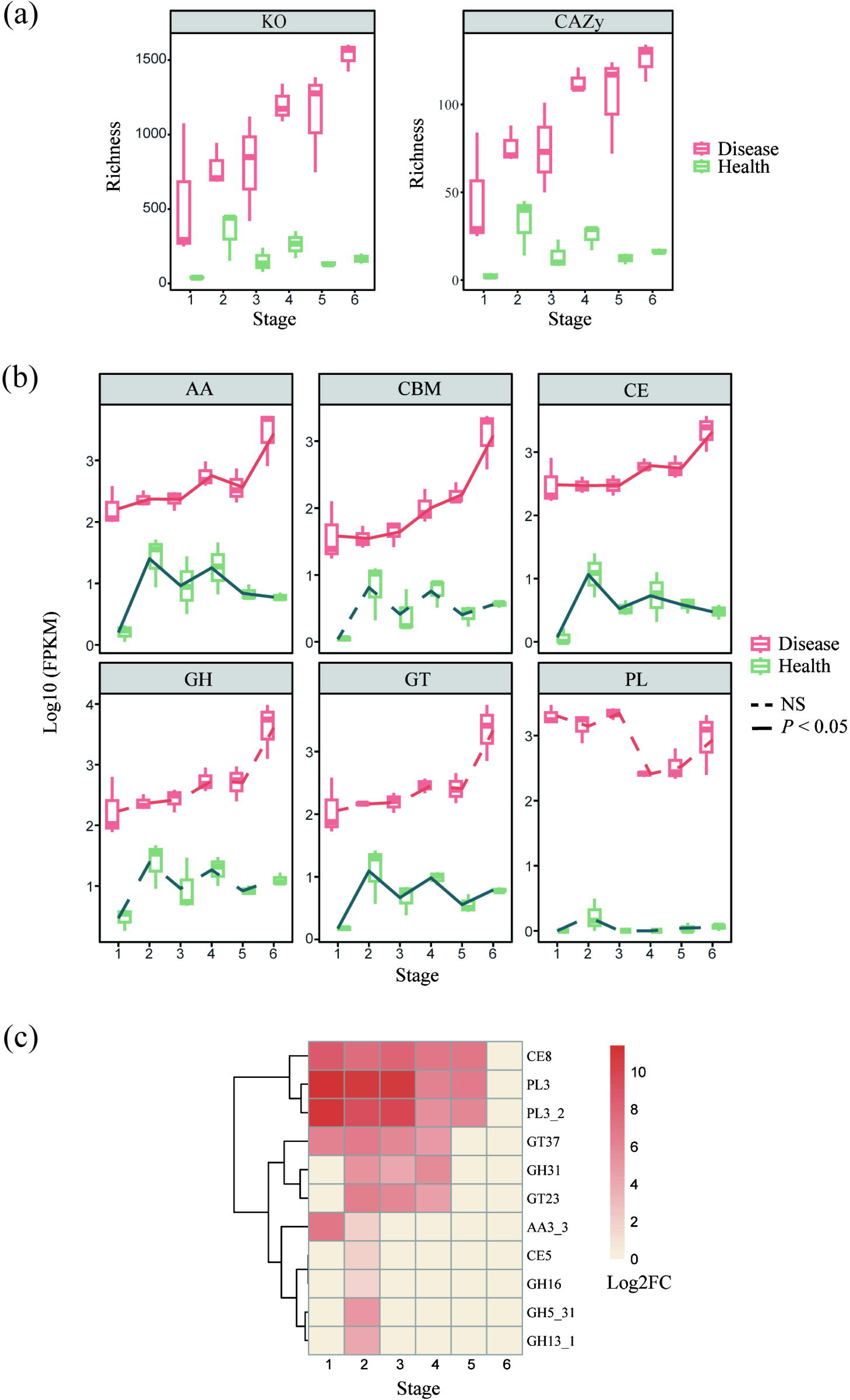

## References

1. Vandenkoornhuyse P, Quaiser A, Duhamel M, Le Van A, Dufresne A. The importance of the microbiome of the plant holobiont. New Phytologist 2015;206(4):1196–206.

2. Simon JC, Marchesi JR, Mougel C, Selosse MA. Host-microbiota interactions: from holobiont theory to analysis. Microbiome 2019;7(1):5.

3. Trivedi P, Leach JE, Tringe SG, Sa T, Singh BK. Plant–microbiome interactions: from community assembly to plant health. Nat Rev Microbiol 2020;18(11):607–21.

4. Berg G, Rybakova D, Fischer D, Cernava T, Vergès MCC, Charles T, et al. Microbiome definition re-visited: old concepts and new challenges. Microbiome 2020;8(1):103.

5. Chialva M, Lanfranco L, Bonfante P. The plant microbiota: composition, functions, and engineering. Current Opinion in Biotechnology 2022;73:135–42.

6. Vayssier-Taussat M, Albina E, Citti C, Cosson JF, Jacques MA, Lebrun MH, et al. Shifting the paradigm from pathogens to pathobiome: new concepts in the light of meta-omics. Front Cell Infect Microbiol 2014;4.

7. Gu S, Wei Z, Shao Z, Friman VP, Cao K, Yang T, et al. Competition for iron drives phytopathogen control by natural rhizosphere microbiomes. Nat Microbiol 2020;5(8):1002–10.

8. Zhang Y, Cao B, Pan Y, Tao S, Zhang N. Metabolite-Mediated Responses of Phyllosphere Microbiota to Rust Infection in Two Malus Species. Microbiology Spectrum 2023;11(2):e03831–22.

9. Zhou X, Zhang J, Khashi u Rahman M, Gao D, Wei Z, Wu F, et al. Interspecific plant interaction via root exudates structures the disease suppressiveness of rhizosphere microbiomes. Molecular Plant 2023;16(5):849–64.

10. Pieterse CMJ, Zamioudis C, Berendsen RL, Weller DM, Van Wees SCM, Bakker PAHM. Induced systemic resistance by beneficial microbes. Annu Rev Phytopathol 2014;52:347–75.

11. Hacquard S, Spaepen S, Garrido-Oter R, Schulze-Lefert P. Interplay Between Innate Immunity and the Plant Microbiota. Annu Rev Phytopathol 2017;55:565–89.

12. Noman M, Ahmed T, Ijaz U, Shahid M, Azizullah, Li D, et al. Plant–Microbiome Crosstalk: Dawning from Composition and Assembly of Microbial Community to Improvement of Disease Resilience in Plants. International Journal of Molecular Sciences 2021;22(13):6852.

13. Pereira LB, Thomazella DPT, Teixeira PJPL. Plant-microbiome crosstalk and disease development. Current Opinion in Plant Biology 2023;72:102351.

14. Kwak MJ, Kong HG, Choi K, Kwon SK, Song JY, Lee J, et al. Rhizosphere microbiome structure alters to enable wilt resistance in tomato. Nat Biotechnol 2018;36(11):1100–9.

15. Berendsen RL, Vismans G, Yu K, Song Y, De Jonge R, Burgman WP, et al. Disease-induced assemblage of a plant-beneficial bacterial consortium. The ISME Journal 2018;12(6):1496–507.

16. Liu H, Li J, Carvalhais LC, Percy CD, Prakash Verma J, Schenk PM, et al. Evidence for the plant recruitment of beneficial microbes to suppress soil-borne pathogens. New Phytologist 2021;229(5):2873–85.

17. Yuan J, Zhao J, Wen T, Zhao M, Li R, Goossens P, et al. Root exudates drive the soil-borne legacy of aboveground pathogen infection. Microbiome 2018;6(1):156.

18. Rolfe SA, Griffiths J, Ton J. Crying out for help with root exudates: adaptive mechanisms by which stressed plants assemble health-promoting soil microbiomes. Current Opinion in Microbiology 2019;49:73–82.

19. Mannaa M, Seo YS. Plants under the Attack of Allies: Moving towards the Plant Pathobiome Paradigm. Plants 2021;10(1):125.

20. Gao M, Xiong C, Gao C, Tsui CKM, Wang MM, Zhou X, et al. Disease-induced changes in plant microbiome assembly and functional adaptation. Microbiome 2021;9(1):187.

21. Carrión VJ, Perez-Jaramillo J, Cordovez V, Tracanna V, de Hollander M, Ruiz-Buck D, et al. Pathogen-induced activation of disease-suppressive functions in the endophytic root microbiome. Science 2019;366(6465):606–12.

22. Bass D, Stentiford GD, Wang HC, Koskella B, Tyler CR. The Pathobiome in Animal and Plant Diseases. Trends in Ecology & Evolution 2019;34(11):996–1008.

23. Lv T, Zhan C, Pan Q, Xu H, Fang H, Wang M, et al. Plant pathogenesis: Toward multidimensional understanding of the microbiome. iMeta 2023;2(3):e129.

24. Yan ZZ, Wang J, Liang J, Batista BD, Liu H, Xiong C, et al. Evidence of distinct response of soil viral community to a plant infection and the disease pathobiome. Journal of Sustainable Agriculture and Environment 2023;2(4):382–7.

25. Lu P, Shi H, Tao J, Jin J, Wang S, Zheng Q, et al. Metagenomic insights into the changes in the rhizosphere microbial community caused by the root-knot nematode *Meloidogyne incognita* in tobacco. Environmental Research 2023;216:114848.

26. Tian BY, Cao Y, Zhang KQ. Metagenomic insights into communities, functions of endophytes and their associates with infection by root-knot nematode, *Meloidogyne incognita*, in tomato roots. Sci Rep 2015;5(1):17087.

27. Malacrinò A, Abdelfattah A, Berg G, Benitez MS, Bennett AE, Böttner L, et al. Exploring microbiomes for plant disease management. Biological Control 2022;169:104890.

28. Vorholt JA. Microbial life in the phyllosphere. Nat Rev Microbiol 2012;10(12):828–40.

29. Chen T, Nomura K, Wang X, Sohrabi R, Xu J, Yao L, et al. A plant genetic network for preventing dysbiosis in the phyllosphere. Nature 2020;580(7805):653–7.

30. Li PD, Zhu ZR, Zhang Y, Xu J, Wang H, Wang Z, et al. The phyllosphere microbiome shifts toward combating melanose pathogen. Microbiome 2022;10(1):56.

31. Liu H, Brettell LE, Singh B. Linking the Phyllosphere Microbiome to Plant Health. Trends in Plant Science 2020;25(9):841–4.

32. Díaz-Cruz GA, Cassone BJ. Changes in the phyllosphere and rhizosphere microbial communities of soybean in the presence of pathogens. FEMS Microbiology Ecology 2022;98(3):fiac022.

33. Seybold H, Demetrowitsch TJ, Hassani MA, Szymczak S, Reim E, Haueisen J, et al. A fungal pathogen induces systemic susceptibility and systemic shifts in wheat metabolome and microbiome composition. Nat Commun 2020;11(1):1910.

34. Tao SQ, Auer L, Morin E, Liang YM, Duplessis S. Transcriptome Analysis of Apple Leaves Infected by the Rust Fungus *Gymnosporangium yamadae* at Two Sporulation Stages. Molecular Plant-Microbe Interactions 2020;33:444–461.

35. Peng D, Wang Z, Tian J, Wang W, Guo S, Dai X, et al. Phyllosphere bacterial community dynamics in response to bacterial wildfire disease: succession and interaction patterns. Front Plant Sci 2024;15.

36. Lemanceau P, Blouin M, Muller D, Moënne-Loccoz Y. Let the Core Microbiota Be Functional. Trends in Plant Science 2017;22(7):583–95.

37. Guo L, Cao Y, Quan J, Liu BY. Crabapple in China: past, present and future. Acta Horticulturae 2019;1263:55–60.

38. Kern FD. A revised taxonomic account of Gymnosporangium. Pennsylvania State University Press, State College. PA 1973.

39. Cummins G, Hiratsuka Y. Illustrated genera of rust fungi. APS Press, St Paul, MN 2003.

40. Yue C, Du C, Wang X, Tan Y, Liu X, Fan H. Powdery mildew-induced changes in phyllosphere microbial community dynamics of cucumber. FEMS Microbiology Ecology 2024;100(5):fiae050.

41. Li Y, Tao S, Liang Y. Time-Course Responses of Apple Leaf Endophytes to the Infection of *Gymnosporangium yamadae*. Journal of Fungi 2024;10(2):128.

42. Collinge DB, Jensen B, Jørgensen HJ. Fungal endophytes in plants and their relationship to plant disease. Current Opinion in Microbiology 2022;69:102177.

43. El-Sheekh MM, Deyab MA, Hasan RSA, Abu Ahmed SE, Elsadany AY. Biological control of *Fusarium* tomato-wilt disease by cyanobacteria *Nostoc* spp. Arch Microbiol 2022;204(1):116.

44. Shiomi HF, Silva HSA, Melo IS de, Nunes FV, Bettiol W. Bioprospecting endophytic bacteria for biological control of coffee leaf rust. Sci agric (Piracicaba, Braz) 2006;63:32–9.

45. Murugaiyan S, Ramasamy K. Isolation and Characterization of Tomato Leaf Phyllosphere *Methylobacterium* and Their Effect on Plant Growth. International Journal of Current Microbiology and Applied Sciences 2017;6:2121–36.

46. Vandepol N, Liber J, Yocca A, Matlock J, Edger P, Bonito G. *Linnemannia elongata* (Mortierellaceae) stimulates *Arabidopsis thaliana* aerial growth and responses to auxin, ethylene, and reactive oxygen species. PLOS ONE 2022;17(4):e0261908.

47. Chen SF, Morgan DP, Michailides TJ. Botryosphaeriaceae and Diaporthaceae associated with panicle and shoot blight of pistachio in California, USA. Fungal Diversity 2014;67(1):157–79.

48. White J, Sullivan R, Moy M, Patel R, Duncan R. An overview of problems in the classification of Plant-parasitic *Clavicipataceae*. Studies in Mycology 2000;45:95–105.

49. Thomma BPHJ, Van Esse HP, Crous PW, De Wit PJGM. *Cladosporium fulvum* (syn. *Passalora fulva*), a highly specialized plant pathogen as a model for functional studies on plant pathogenic Mycosphaerellaceae. Molecular Plant Pathology 2005;6(4):379–93.

50. Faust K, Raes J. Microbial interactions: from networks to models. Nat Rev Microbiol 2012;10(8):538–50.

51. Wei Z, Gu Y, Friman VP, Kowalchuk GA, Xu Y, Shen Q, et al. Initial soil microbiome composition and functioning predetermine future plant health. Science Advances 2019;5(9):eaaw0759.

52. Das S, Rabha J, Narzary D. Assessment of soil yeasts *Papiliotrema laurentii* S-08 and *Saitozyma podzolica* S-77 for plant growth promotion and biocontrol of *Fusarium* wilt of brinjal. Journal of Applied Microbiology 2023;134(11):lxad252.

53. Li F, Chen L, Redmile-Gordon M, Zhang J, Zhang C, Ning Q, et al. *Mortierella Elongata*’s Roles in Organic Agriculture and Crop Growth Promotion in a Mineral Soil. Land Degradation & Development 2018;29.

54. Ozimek E, Hanaka A. Mortierella Species as the Plant Growth-Promoting Fungi Present in the Agricultural Soils. Agriculture 2021;11(1):7.

55. Louca S, Polz MF, Mazel F, Albright MBN, Huber JA, O’Connor MI, et al. Function and functional redundancy in microbial systems. Nat Ecol Evol 2018;2(6):936–43.

56. Paasch BC, He SY. Toward understanding microbiota homeostasis in the plant kingdom. PLOS Pathogens 2021;17(4):e1009472.

57. Chung KR. Mitogen-activated protein kinase signaling pathways of the tangerine pathotype of *Alternaria alternata*. MAP Kinase 2013;2:e4.

58. Chen S, Zhou Y, Chen Y, Gu J. fastp: an ultra-fast all-in-one FASTQ preprocessor. Bioinformatics 2018;34(17):i884–90.

59. Li H, Durbin R. Fast and accurate short read alignment with Burrows-Wheeler transform. Bioinformatics 2009;25(14):1754–60.

60. Kopylova E, Noé L, Touzet H. SortMeRNA: fast and accurate filtering of ribosomal RNAs in metatranscriptomic data. Bioinformatics 2012;28(24):3211–7.

61. Haas BJ, Papanicolaou A, Yassour M, Grabherr M, Blood PD, Bowden J, et al. De novo transcript sequence reconstruction from RNA-seq using the Trinity platform for reference generation and analysis. Nat Protoc 2013;8(8):1494–512.

62. Fu L, Niu B, Zhu Z, Wu S, Li W. CD-HIT: accelerated for clustering the next-generation sequencing data. Bioinformatics 2012;28(23):3150–2.

63. Buchfink B, Xie C, Huson DH. Fast and sensitive protein alignment using DIAMOND. Nat Methods 2015;12(1):59–60.

64. Li B, Dewey CN. RSEM: accurate transcript quantification from RNA-Seq data with or without a reference genome. BMC Bioinformatics 2011;12:323.

65. Oksanen J, Kindt R, Legendre P, O’Hara B, Stevens MHH, Oksanen MJ, Suggests M. The vegan package. Community Ecology Package 2007;10: 631–637.

66. Hothorn T, Bretz F, Westfall P. Simultaneous Inference in General Parametric Models. Biometrical Journal 2008;50(3):346–63.

67. Aphalo PJ, Slowikowski K, Mouksassi S. ggpmisc: Miscellaneous Extensions to ‘ggplot2’ 2024.

68. Gehlenborg N. UpSetR: A More Scalable Alternative to Venn and Euler Diagrams for Visualizing Intersecting Sets. 2015.

69. Love MI, Huber W, Anders S. Moderated estimation of fold change and dispersion for RNA-seq data with DESeq2. Genome Biol 2014;15(12):550.

70. Gu L. TCseq: time course sequencing data analysis. 2023.

71. Potter SC, Luciani A, Eddy SR, Park Y, Lopez R, Finn RD. HMMER web server: 2018 update. Nucleic Acids Research 2018;46(W1):W200–4.

72. Wu T, Hu E, Xu S, Chen M, Guo P, Dai Z, et al. clusterProfiler 4.0: A universal enrichment tool for interpreting omics data. Innovation (Camb) 2021;2(3):100141.

73. Harrell Jr. Hmisc: Harrell Miscellaneous version 5.1-3 from CRAN. 2024

74. Bastian M, Heymann S, Jacomy M. Gephi: An Open Source Software for Exploring and Manipulating Networks. Proceedings of the International AAAI Conference on Web and Social Media 2009;3(1):361–2.

75. Archer E. rfPermute: Estimate Permutation p-Values for Random Forest Importance Metrics. 2011.

